# Intact and Degenerate Diguanylate Cyclases regulate *Shigella* Cyclic di-GMP

**DOI:** 10.1101/2024.04.08.588579

**Authors:** Ruchi Ojha, Stefanie Krug, Prentiss Jones, Benjamin J. Koestler

## Abstract

The intracellular human pathogen *Shigella* invades the colonic epithelium to cause disease. Prior to invasion, this bacterium navigates through different environments within the human body, including the stomach and the small intestine. To adapt to changing environments, *Shigella* uses the bacterial second messenger c-di-GMP signaling system, synthesized by diguanylate cyclases (DGCs) encoding GGDEF domains. *Shigella flexneri* encodes a total of 9 GGDEF or GGDEF-EAL domain enzymes in its genome, but 5 of these genes have acquired mutations that presumably inactivated the c-di-GMP synthesis activity of these enzymes. In this study, we examined individual *S. flexneri* DGCs for their role in c-di-GMP synthesis and pathogenesis. We individually expressed each of the 4 intact DGCs in an *S. flexneri* strain where these 4 DGCs had been deleted (Δ4DGC). We found that the 4 *S. flexneri* intact DGCs synthesize c-di-GMP at different levels *in vitro* and during infection of tissue-cultured cells. We also found that *dgcF* and *dgcI* expression significantly reduces invasion and plaque formation, and *dgcF* expression increases acid sensitivity, and that these phenotypes did not correspond with measured c-di-GMP levels. However, deletion of these 4 DGCs did not eliminate *S. flexneri* c-di-GMP, and we found that *dgcE, dgcQ,* and *dgcN*, which all have nonsense mutations prior to the GGDEF domain, still produce c-di-GMP. These *S. flexneri* degenerate DGC genes are expressed as multiple proteins, consistent with multiple start codons within the gene. We propose that both intact and degenerate DGCs contribute to *S. flexneri* c-di-GMP signaling.

## Introduction

Shigellosis, caused by the human pathogen *Shigella,* is a prominent gastrointestinal infection in developing countries (1, 2). *Shigella flexneri* evolved from commensal *E. coli* to efficiently infect the human gastrointestinal tract after navigating different host microenvironments (3, 4). With the global rise in antibiotic resistance among *Shigella* strains (5, 6), and the lack of vaccines for Shigellosis prevention (1, 7), it is imperative to understand *Shigella* pathogenesis mechanisms. In the colon, *Shigella* invades the epithelium using a type III secretion system (T3SS) encoded on a ∼220kb virulence plasmid (7–10). Once inside colonic epithelial cells, *Shigella* multiplies and spreads using actin-based motility (11–13).

During infection, *S. flexneri* senses and responds to many different environmental signals to adapt to various microenvironments within the human body. For example, lower pH in the stomach alters the expression of acid-related genes and induces biofilm formation (14, 15), bile acids in the small intestine promote initial adhesion and invasion (16–18), and formate within a host cell promotes cell-to-cell spread (19). Bacteria like *S. flexneri* encode many different systems to sense and respond to these signals, one of which is the second messenger cyclic-di-guanosine monophosphate (c-di-GMP) signaling system (20, 21).

C-di-GMP signaling is widely conserved in most bacteria (22, 23) and regulates diverse phenotypes (24–28). C-di-GMP synthesis is driven by diguanylate cyclases (DGCs) encoding a C-terminal GGDEF domain, which brings together two GTP molecules. Alterations of the GGDEF catalytic site can eliminate c-di-GMP synthesis of these enzymes (29–31). Conversely, c-di-GMP is degraded by c-di-GMP specific phosphodiesterases (PDEs) encoding an EAL domain (22, 23, 28). In many bacteria, c-di-GMP promotes biofilm formation and reduces invasive capacity (7, 20, 32). However, certain DGCs and PDEs synthesize local pools of c-di-GMP, allowing them to regulate specific phenotypes regardless of overall c-di-GMP levels; this phenomenon is known as signaling specificity (31, 33, 34). *E. coli* K12 encodes 19 GGDEF-containing genes in its genome (35); in comparison, *S. flexneri* encodes 9 GGDEF or GGDEF-EAL containing genes, but the majority of these genes contain mutations that presumably eliminate c-di-GMP synthesis activity (pseudogenes) (36). *S. flexneri* has four DGCs predicted to synthesize c-di-GMP: *dgcC*, *dgcF*, *dgcI,* and *dgcP* (36).

We previously showed that deletion of intact *S. flexneri* DGCs alters pathogenesis-related phenotypes (20). Here, we investigate how *S. flexneri* DGCs contribute to c-di-GMP levels and how the expression of these genes impacts different phenotypes. To this end, we created a mutant *S. flexneri* strain where all 4 intact DGCs were deleted (Δ4DGC), and then expressed each *S. flexneri* DGC and respective active site mutant DGCs (GG➔AA) from plasmids. We observed that all 4 DGCs were able to synthesize c-di-GMP at different levels and that each gene differentially contributed to acid resistance, invasion, and plaque formation. Interestingly, we still detected c-di-GMP in our *S. flexneri* Δ4DGC strain, which led us to examine the 5 putative *S. flexneri* DGC pseudogenes. We found that expression of *S. flexneri dgcE, dgcQ* and *dgcN*, each of which have nonsense mutations prior to the GGDEF domain, increased c-di-GMP, and deletion of *S. flexneri* Δ*dgcE* (Δ5DGC) and Δ*dgcE*Δ*dgcQ* (Δ6DGC) from the Δ4DGC strain significantly reduced c-di-GMP. This provides evidence that *S. flexneri* pseudogenes can retain function.

## Results

### *S. flexneri* DGCs increase c-di-GMP

The *S. flexneri* 2457T genome encodes four intact GGDEF domain enzymes, *dgcC* (S0329), *dgcF* (S1698), *dgcI* (S0827) and *dgcP* (S1545) (36). Individual deletion of DGCs from the *S. flexneri* genome significantly reduces biofilm formation and alters invasion, plaque formation and acid resistance in a DGC-specific manner (20). To study how individual *S. flexneri* DGCs contribute to pathogenesis, we generated a mutant strain where all four intact DGCs (*dgcC*, *dgcF*, *dgcI* and *dgcP*) were deleted (Δ4DGC). Each of these four *S. flexneri* DGCs were then individually expressed from an isopropyl-b-D-thiogalactopyranoside (IPTG)-inducible plasmid in the *S. flexneri* Δ4DGC background. As a negative control, we mutated two amino acids from the active site of each of the four *S. flexneri* DGCs from GG(D/E)EF to AA(D/E)EF (Glycine to Alanine; GG→AA) to separate the regulatory role of the protein itself to its c-di-GMP synthesis (29). This mutation has been shown to eliminate c-di-GMP synthesis in DGCs from other bacteria (29, 30, 37). We confirmed thathe t expression of these 4 DGCs did not alter *S. flexneri* growth in broth (Fig. S1A).

Ectopic expression of the *V. cholerae* DGC VCA0956 significantly increases c-di-GMP in *S. flexneri* as measured by liquid chromatography coupled with mass spectrometry (LC-MS/MS) (20). We hypothesized that the expression of intact *S. flexneri* DGCs will increase c-di-GMP levels as compared to GG→AA mutant alleles. C-di-GMP from these strains was extracted during mid-log growth (∼3 hours post IPTG addition) and quantified using LC-MS/MS. We observed that *dgcI* expression resulted in the highest levels of c-di-GMP, whereas *dgcC*, and *dgcF* expression increased c-di-GMP to levels comparable to the VCA0956 expression strain. Contrary to that, the *dgcP* expression strain showed no difference in c-di-GMP levels to that of the Δ4DGC strain (Fig. 1). C-di-GMP was below our limit of detection when individual GG→AA DGC mutants were expressed, but interestingly, we observed detectable levels of c-di-GMP in the *S. flexneri* Δ4DGC strain carrying an empty plasmid, comparable to the WT strain (Fig.1).

**Figure 1.**
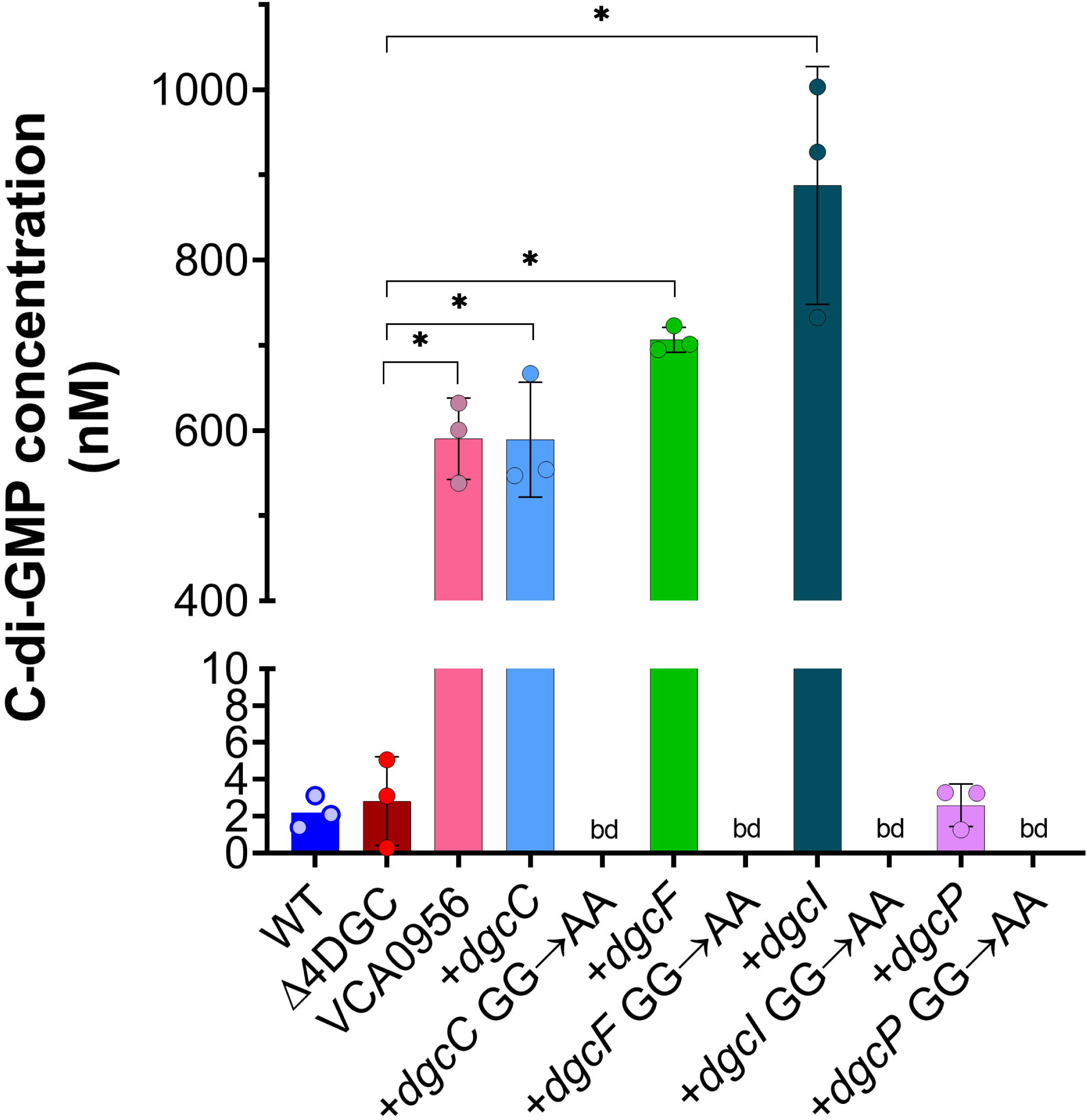
*S. flexneri* intact DGCs regulate c-di-GMP levels. Individual *S. flexneri* DGCs were expressed in the *S. flexneri* Δ4DGC strain, and c-di-GMP levels were measured using LC-MS/MS. *dgcC*, *dgcF* and *dgcI* expression in the Δ4DGC strain showed significantly more c-di-GMP synthesis than the GG→AA mutants or the empty vector controls, extracted at mid-log phase. Expression of *dgcP* did not significantly alter *S. flexneri* c-di-GMP levels in comparison to the Δ4DGC strain. We also noticed no significant difference between WT *S. flexneri* and the Δ4DGC strain. VCA0956 (positive control) significantly increased c-di-GMP levels in *S. flexneri* (20). bd indicates below detection, * represents significant differences between strains and the Δ4DGC strain carrying an empty vector, as analyzed by one-way ANOVA with Sidak’s multiple comparisons posttest (*p<0.05*). Each symbol represents independent replicates, error bars indicate standard deviation.

### *S. flexneri dgcP* expression increases c-di-GMP 3 hours post-induction

We were intrigued that *S. flexneri dgcP* expression did not increase c-di-GMP levels compared to the other three DGCs (Fig. 1). We further wanted to investigate temporal dynamics of the four intact *S. flexneri* DGCs; thus, as a complementary approach, we quantified relative c-di-GMP levels using a plasmid encoding a double tandem riboswitch Bc3-Bc4 controlling the expression of the reporter gene m-Scarlett I (riboswitch reporter) (38), allowing c-di-GMP measurement with single-cell resolution of live cells by microscopy. The riboswitch reporter binds c-di-GMP with high affinity and specificity, and in the absence of c-di-GMP, a terminator inhibits the translation of m-Scarlett I. This c-di-GMP reporter was validated in the closely related *E. coli* (38).

We hypothesized that *S. flexneri* DGC expression would result in m-Scarlett I fluorescence that correlates with c-di-GMP levels measured by LC-MS/MS. We quantified m-Scarlett I fluorescence of our *S. flexneri* DGC expression strains over time using live-cell fluorescence microscopy. We then compared the mean of individual fluorescent cells of each DGCs to the GG→AA mutants. Of note, we observed no differences in cell size and growth among all our strains. As expected, c-di-GMP-dependent fluorescence increased rapidly and significantly over time in *S. flexneri dgcC*, *dgcF*, and *dgcI*, reaching near maximal fluorescence after 1 hour (Fig. 2A-C). Interestingly, *S. flexneri dgcP* expression increased fluorescence at a slower rate, resulting in significantly higher c-di-GMP levels 3 hours post IPTG induction compared to the GG→AA *dgcP* mutant (Fig. 2D). We observed similar patterns when the fluorescence of these strains was quantified using a plate reader (Fig. S2). We speculate that differences in the growth conditions used for quantitation by LC-MS/MS and microscopy alter *S. flexneri* DgcP c-di-GMP synthesis (Fig 1 and 2D).

**Figure 2.**
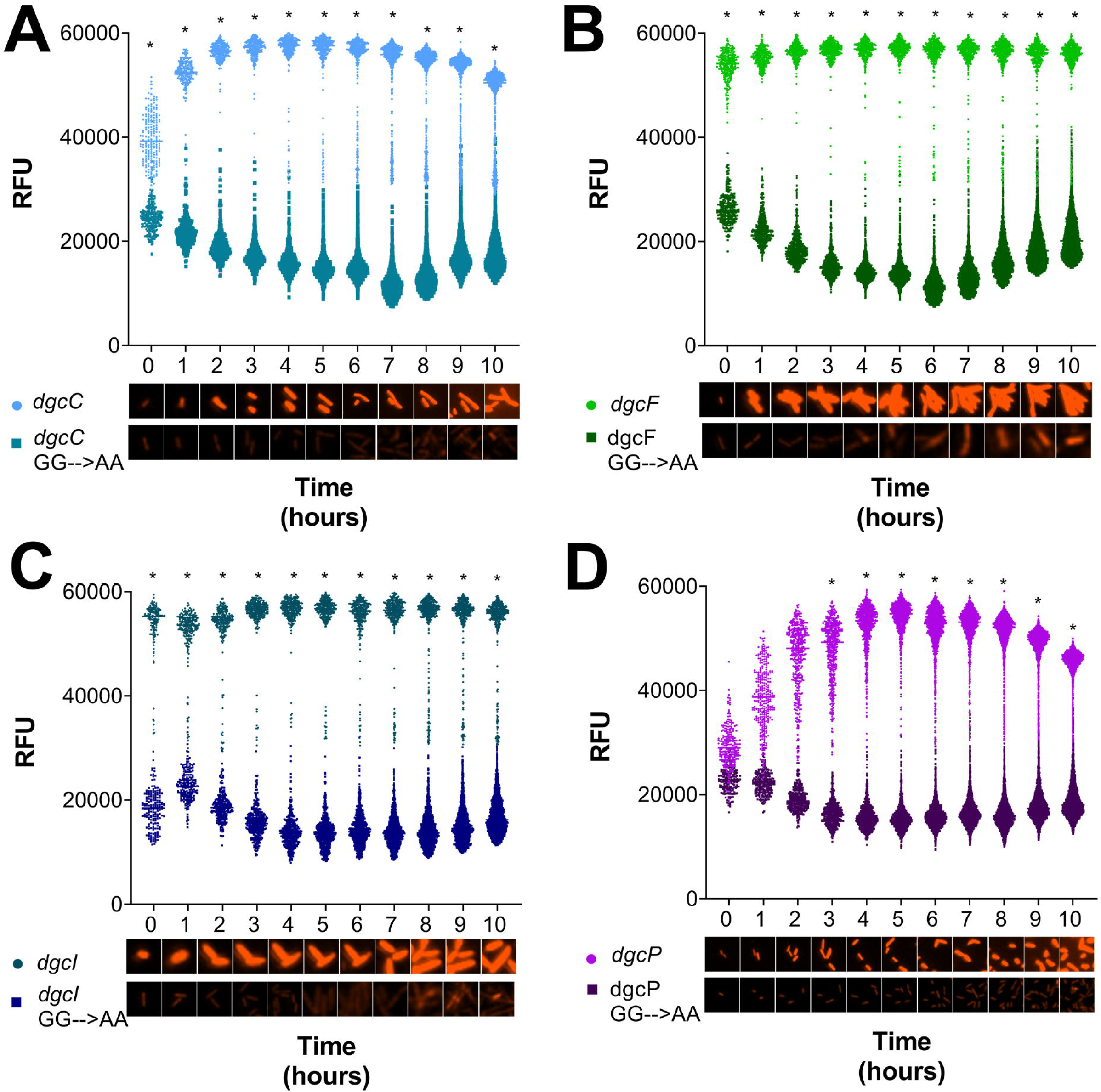
DGC expression significantly increases c-di-GMP-dependent fluorescence compared to GG→AA mutants. Single bacterial cell fluorescence (relative fluorescence units, RFU) was measured using microscopy over time for the *S. flexneri* Δ4DGC strain expressing (A) *dgcC*, (B) *dgcF*, (C) *dgcI*, and (D) *dgcP*, alongside their respective GG→AA mutants. *S. flexneri* DGCs were expressed from an IPTG-inducible plasmid, and a second plasmid expressing the c-di-GMP-specific double tandem riboswitch with m-Scarlett I (riboswitch reporter) was used to measure c-di-GMP-dependent fluorescence. Each of the four DGCs fluoresce more than the GG→AA mutant DGCs. Symbols (circles: DGCs; squares: GG→AA alleles) indicate individual fluorescent cells for each time point for each strain. Graph is representative of one of the three independent trials. Micrographs below (A-D) are representative samples of individual bacterial cells; red indicates c-di-GMP-dependent m-Scarlett I fluorescence. Significant differences in fluorescence between DGCs and GG→AA alleles were analyzed using two-way ANOVA with Sidak’s multiple comparisons posttest (*, *p<0.05*) at each time point.

### *S. flexneri* c-di-GMP levels are dynamic during host-cell growth

As *S. flexneri* is an intracellular pathogen, we were interested in c-di-GMP changes during growth in a host cell, and if the 4 intact *S. flexneri* DGCs are responsible for c-di-GMP production in this environment. To answer these questions, we used our riboswitch reporter to evaluate changes in c-di-GMP production of live *S. flexneri* cells over time. When WT *S. flexneri* was grown in a defined medium, c-di-GMP remained relatively constant, exhibiting modestly decreasing fluorescence over time, although this trend was not statistically significant. The *S. flexneri* Δ4DGC strain was consistently lower than the WT at all time points, although this difference too was not statistically significant (Fig 3A).

**Figure 3.**
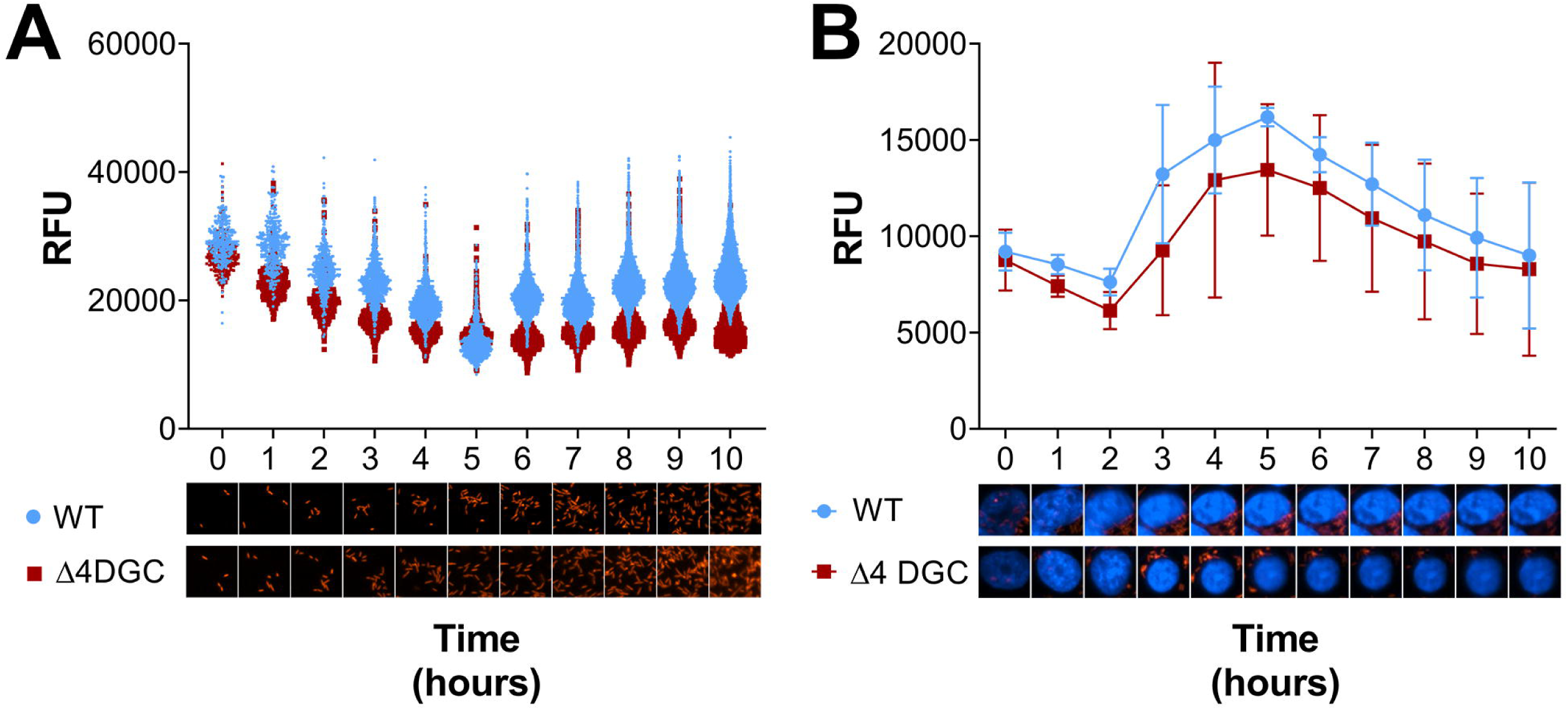
*S. flexneri* increases c-di-GMP production in vivo 3 hours post infection. C-di-GMP-driven fluorescence (RFU) of *S. flexneri* WT and the Δ4DGC strain carrying the c-di-GMP riboswitch reporter grown (A) in culture or (B) in Henle-407 cells. (A) In culture, c-di-GMP levels remained relatively constant over the course of 10 hours, and were consistently but non-significantly higher in the WT *S. flexneri* strain than the Δ4DGC strain. Symbols indicate the individual fluorescent cells, representative of one trial among three. (B) In both the *S. flexneri* WT and Δ4DGC strain, c-di-GMP levels increased 3 HPI and then gradually decreased starting at 5 HPI. Micrographs below are representative of individual bacterial cells in culture. Infected Henle-407 cells were stained with Hoechst to label host cell nuclei (blue). Red indicates c-di-GMP dependent m-Scarlett I fluorescence. Significant differences between the WT and Δ4DGC strain were analyzed using two-way ANOVA with Sidak’s multiple comparisons posttest (*, *p<0.05*); no statistical significance was observed.

We performed the same analysis to assess changing c-di-GMP levels over time during *S. flexneri* infection of Henle-407 cells (Fig. 3B) (39). In contrast to *S. flexneri* grown in culture, WT *S. flexneri* fluorescence increased at 3 hours post infection (HPI), peaked 5 HPI, and then gradually decreased fluorescence during Henle-407 infection. For the *S. flexneri* Δ4DGC strain, we again observed that c-di-GMP levels were consistently non-significantly lower than in the WT strain, but we saw the same general pattern, where fluorescence increased at 3 HPI, peaked at 5 HPI, and then decreased afterwards (Fig 3B).

### *S. flexneri* DGCs produce c-di-GMP during host cell infection

To determine how *S. flexneri* DGCs contribute to changing c-di-GMP levels during host cell growth, we compared the expression of *S. flexneri dgcC*, *dgcF*, *dgcI*, *dgcP* and their respective GG→AA mutants in the *S. flexneri* Δ4DGC strain during infection. Expression of each of the 4 intact DGCs increased c-di-GMP, but the temporal dynamics were different for each strain (Fig. 4). Specifically, the time that it took each DGC to initiate c-di-GMP synthesis varied. *dgcC* expression exhibited no significant fluorescence until 3 HPI, and then increased rapidly (Fig. 4A). *dgcF* expression resulted in constitutively high c-di-GMP throughout the entire course of infection (Fig. 4B). *dgcI* expression did not increase fluorescence until 2 HPI (Fig. 4C). Similar to *dgcI*, *dgcP* expression increased starting at 2 HPI and peaked at 5 HPI, and then steadily decreased for the remainder of the experiment (Fig. 4D). Interestingly, for each of our GG→AA mutants, we observed an increase in fluorescence that peaked between 4-6 HPI, and then decreased as the infection progressed.

**Figure 4.**
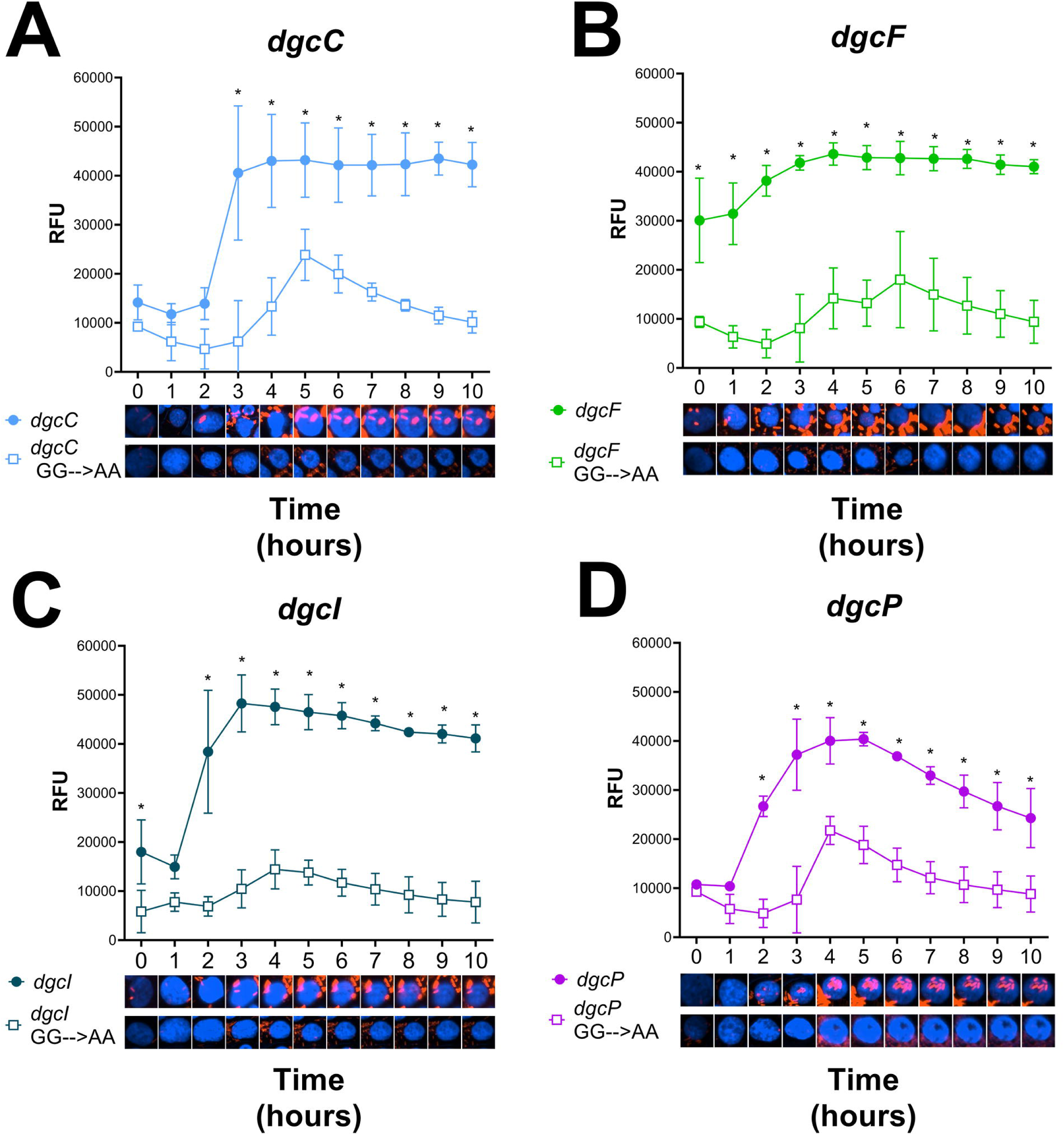
*S. flexneri* DGCs regulate c-di-GMP synthesis post infection. C-di-GMP-dependent fluorescence (RFU) of the *S. flexneri* Δ4DGC strain expressing (A) *dgcC*, (B) *dgcF*, (C) *dgcI*, or (D) *dgcP*, alongside their respective GG→AA mutant alleles, was quantified during infection of Henle-407 cells. Expression of all four intact DGCs significantly increased fluorescence compared to individual GG→AA mutant DGCs, but the time it took for these differences to become significant was variable. Representative micrographs below illustrate single cell fluorescence; red indicates c-di-GMP dependent m-Scarlett I fluorescence, and blue host cell nuclei stained with Hoechst. Symbols indicate the mean fluorescence of single cell measurements from three independent replicates, and error bars indicate standard deviation. Significant differences between strains at each time point were analyzed using two-way ANOVA with Sidak’s multiple comparisons posttest (*, *p<0.05*).

### *dgcF* expression increases *S. flexneri* acid sensitivity in AR1-inducing conditions

During human infection, *Shigella* must survive the highly acidic environment in the stomach before it can access the human colon. *Shigella* survives acid stress with at least two systems (AR1 and AR2) that are induced in different conditions (40–44). We previously showed that deletion of *S. flexneri dgcF* increases acid resistance (20); therefore, we hypothesized that expression of *S. flexneri dgcF* will decrease acid resistance. We pre-conditioned *S. flexneri* DGCs in complex media at pH 5.5 to induce AR1, and separately at pH 5.5 with glucose and low oxygen to induce AR2. We then challenged our DGC expression strains with exposure to defined media at pH 2.5 (to study AR1) or pH 2.5 with glutamate (to study AR2) (40–42, 45). *dgcF* expression significantly reduced acid survival in AR1-inducing conditions, compared to other DGCs and the GG→AA mutants (Fig 5A). In AR2-inducing conditions, DGC expression did not significantly decrease acid resistance (Fig 5B). This suggests that *S. flexneri dgcF* specifically regulates acid survival in AR1-inducing conditions.

**Figure 5.**
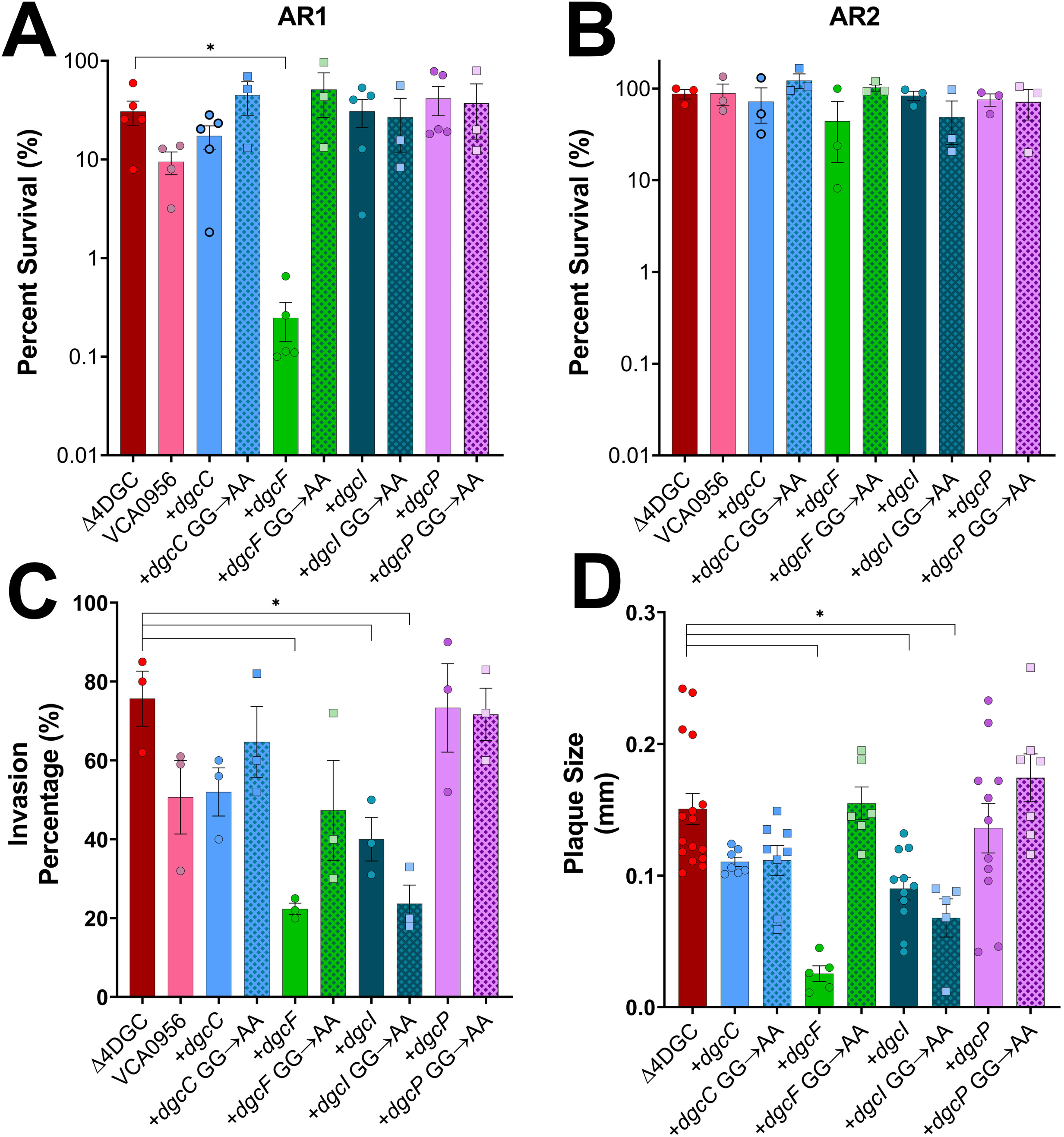
Expression of *S. flexneri dgcF* increases acid sensitivity in AR1-inducing conditions and reduces virulence. (A) Each *S. flexneri* intact DGC was expressed in the Δ4DGC strain and challenged in pH 2.5 medium in AR1-inducing conditions for one hour. Percent survival was determined by quantifying the subsequent CFU/ml of cultures challenged in pH 2.5 medium, compared to organisms in pH 7.0 medium. *dgcF* expression significantly decreased acid resistance as compared to the *S. flexneri* Δ4DGC strain carrying an empty plasmid and the GG→AA mutant *dgcF*, as determined by a Kruskal-Wallis test with Dunn’s multiple comparisons post-test. (B) The same experiment was performed in AR2-inducing conditions, which relies on external glutamate (41, 42). AR2-inducing conditions resulted in overall higher *S. flexneri* survival, and we observed no significant differences in survival when each intact DGC or GG→AA mutant was expressed. Bars represent the mean of three independent trials, and error represents standard deviation. (C) We assessed the capacity of *S. flexneri* strains to invade Henle-407 cells. *dgcF, dgcI* and the *dgcI* GG→AA mutant expression in the Δ4DGC strain show significant reduction in the percentage of invaded cells. Bars indicate the mean of 3 independent trials, and error represents standard deviation. (D) Plaque size was measured after infecting Henle-407 cells with the *S. flexneri* Δ4DGC strain expressing the 4 intact DGCs or GG→AA mutant alleles. *dgcF*, *dgcI* and the GG→AA mutant *dgcI* strains show significant reduction in plaque size as compared to the *S. flexneri* Δ4DGC mutant strain carrying an empty plasmid. Symbols represent individual plaques from one replicate among the three individual trials. Bars indicate the mean and error bars represents the standard deviation. Significant differences were analyzed using two-way ANOVA with Sidak’s multiple comparisons posttest (*, *p<0.05*).

### Virulence phenotypes in *S. flexneri* are regulated by *dgcF and dgcI*

VCA0956 expression reduces *S. flexneri* invasion and plaque formation, but the *S. flexneri* Δ*dgcF* strain also has reduced invasion and forms smaller plaques in Henle-407 monolayers (20). To see how expressing the 4 intact *S. flexneri* DGCs impacts these phenotypes, we examined the invasion and plaque phenotypes affected by our DGC expression strains. Henle-407 cells were infected with the *S. flexneri* Δ4DGC strain with DGC expression plasmids and their respective GG→AA mutants. As a control, we included VCA0956 expression which decreases invasion (20). Expression of *dgcF* and *dgcI* exhibited invasion defects as compared to the empty plasmid control, while *dgcC* and *dgcP* expression strains show no significant difference in invasion (Fig 5C). Interestingly, the *dgcI* GG→AA mutant also exhibited an invasion defect, while all other GG→AA DGC mutants did not.

We also examined the ability of each DGC expression strain to form plaques in cell culture monolayers, as previously described (39). Each strain was induced 30 minutes post-infection to ensure comparable invasion. Similar to the invasion assay, we found that the *dgcF, dgcI* and *dgcI* GG→AA mutant showed reduced plaque size in comparison to the empty plasmid control, while *dgcC* and *dgcP* expression showed no significant difference in plaque size compared to the Δ4DGC (Fig 5D).

### Four *S. flexneri* DGC pseudogenes encode GGDEF domains after nonsense mutations

Our LC-MS/MS measurements and riboswitch reporter data indicate that the *S. flexneri* Δ4DGC strain still produces c-di-GMP; therefore, we sought to identify other *S. flexneri* genes that synthesize c-di-GMP. We started by investigating the other DGC pseudogenes. In comparison to *E.coli* K12 substr. MG1655 (Accession: NC_000913), 4 GGDEF domain and 1 GGDEF-EAL domain genes are annotated as pseudogenes (considered degenerate) in *S. flexneri* 2457T (Accession: AE014073.1), and include deletions, frameshift mutations, and/or nonsense mutations that presumably disrupt the GGDEF domain (20, 36). Compared to *E. coli*, there are 9 other DGCs that either do not contain the GGDEF domain or are deleted in *S. flexneri* (36). Therefore, for the scope of this study, we focused on the 5 GGDEF and GGDEF-EAL domain encoding genes with premature stop codons (Table 1). Recent studies indicate that other *S. flexneri* degenerate enzymes have demonstrable phenotypes (46); therefore, we hypothesized that one or more of these DGC pseudogenes retain c-di-GMP synthesis. Compared to *E. coli* MG1655, *dgcT*, *dgcZ* and *cdgI* genes are completely absent from the *S. flexneri* 2457T genome (Table 1). *S. flexneri dgcO* (S1870) and *dgcJ* (S1553) genes have truncating deletions that remove the GGDEF domain. The remaining four *S. flexneri* genes, *dgcM* (S1445), *dgcN* (S2841), *dgcE* (S2255) and *dgcQ* (S2095), have pairwise identity >97.9% compared to *E. coli* K12 strain; however, each of these genes encodes a nonsense mutation resulting in a stop codon preceding the GGDEF domain, which we confirmed by whole genome sequencing. Notably, these four DGC genes have alternative reading frames that begin close to the nonsense mutation site and encode an intact GG[D/E]EF domain. *S. flexneri dgcM*, *dgcE* and *dgcQ* all contain an RXXD inhibition site, potentially enabling allosteric feedback inhibition (Table 1) (47, 48). Interestingly, while the 4 intact *S. flexneri* 2475T DGCs are the most well conserved in other *Shigella* spp., degenerate DGCs are intact in the genomes of other *Shigella* spp. (Fig S3).

**Table 1.**
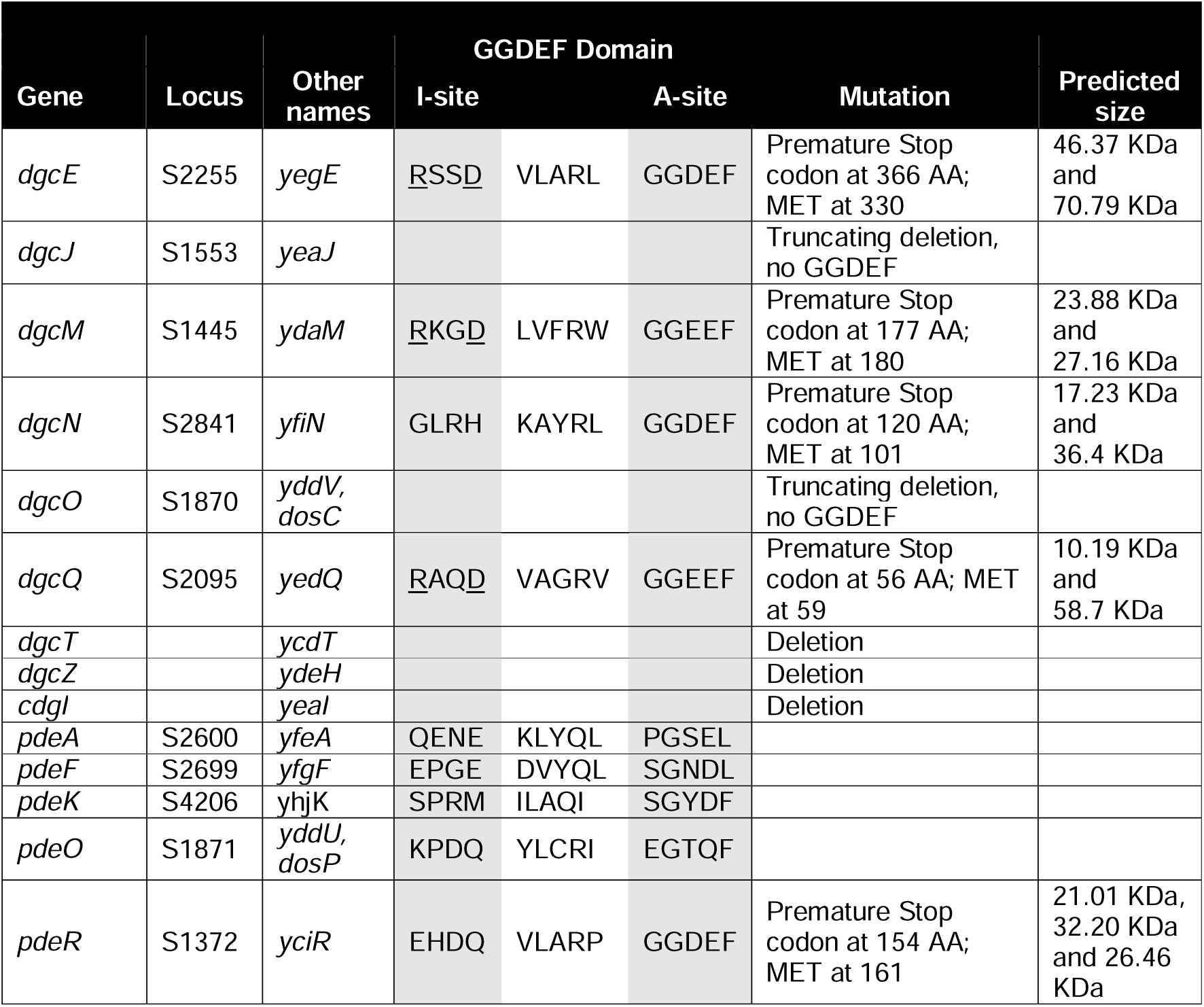
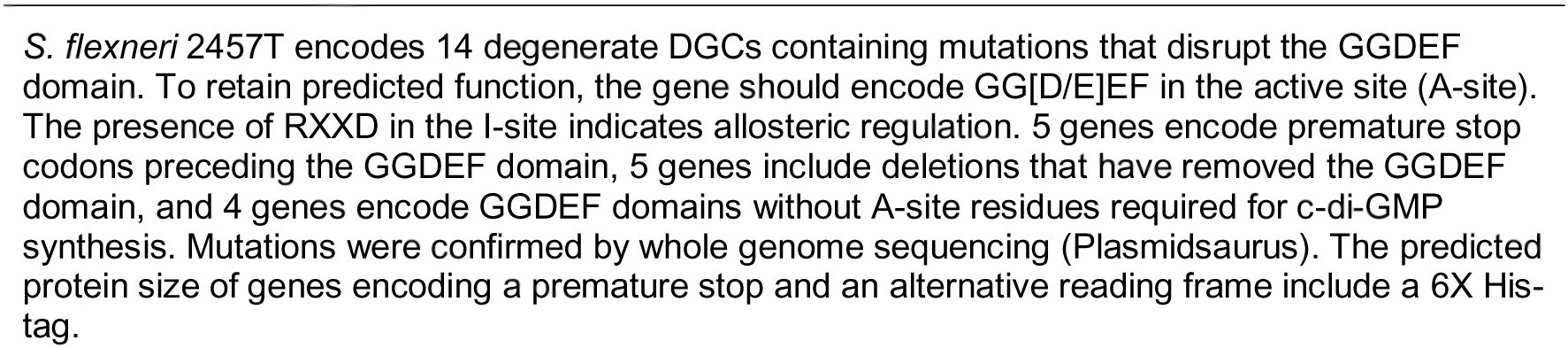
Shigella degenerate and deleted DGCs.

### S. flexneri dgcE and dgcQ produce c-di-GMP

To determine if *S. flexneri* degenerate DGCs are capable of c-di-GMP synthesis, we focused on the five degenerate DGCs that contain nonsense mutations preceding the GGDEF domain. We also included *dgcO* and *dgcJ*, with the expectation that these genes could not produce c-di-GMP due to the complete lack of GGDEF domain sequence. We expressed each of these 7 genes from a pET28a (+) plasmid with 6X His-tag on both N- and C-terminus in *E. coli* BL21 and assessed c-di-GMP levels using our c-di-GMP riboswitch reporter (Fig. 6A). Surprisingly, *dgcE*, *dgcQ* and *dgcN* expression resulted in fluorescence significantly higher than the empty plasmid control and other degenerate DGCs at 5 hours post IPTG induction, consistent with c-di-GMP synthesis (Fig 6A). *dgcM* and *pdeR* expression reduced fluorescence compared to the empty vector control. Expectedly, *dgcO* and *dgcJ* expression did not result in any increase in fluorescence relative to the empty plasmid control.

**Figure 6.**
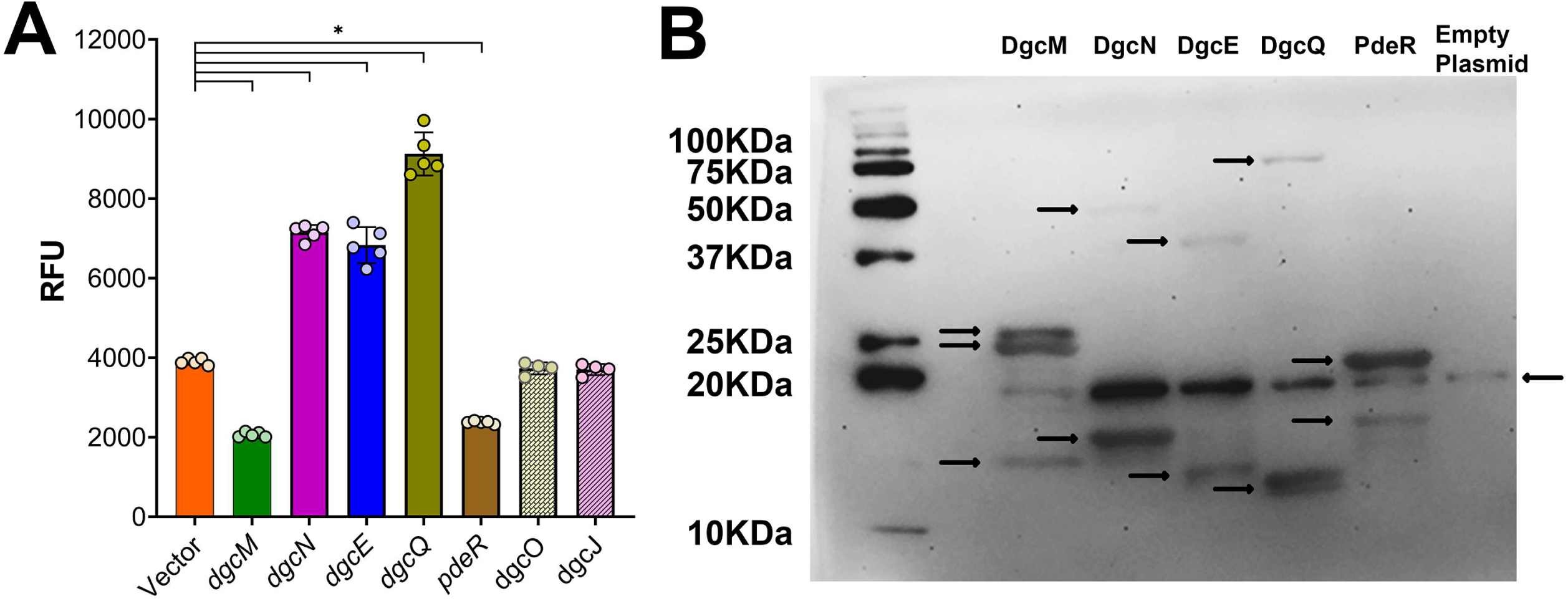
*S. flexneri dgcN, dgcE* and *dgcQ* significantly increase c-di-GMP levels. (A) *S. flexneri* degenerate DGCs were expressed in *E. coli* BL21 and c-di-GMP levels measured using the c-di-GMP riboswitch reporter. *dgcE*, *dgcQ*, and *dgcN* had significantly increased fluorescence in comparison to our empty plasmid control, whereas *dgcM* and *pdeR* showed significantly lower fluorescence. Individual symbols represent three independent replicates, * represents significant differences in comparison to the empty plasmid control as determined by one-way ANOVA with Sidak’s multiple comparisons posttest (p<0.05). Error bars indicate standard deviation. (B) *S. flexneri* degenerate DGCs with both N- and C-terminal 6X His-tags were expressed in *E. coli* BL21, and protein expression was visualized by SDS-PAGE followed by Western blot. Samples for DgcN and DgcR were loaded 2x to visualize faint bands. Arrows highlight fragments for each gene expressed. We observed a non-specific band at ∼20KDa in all lanes including our empty vector control.

We next assessed the protein expression of these 5 degenerate DGCs by Western Blot, using the same plasmids with both N- and C-terminal 6X His-tags. Of note, a non-specific band was observed at ∼20KDa in all samples including our empty vector control (Fig 6B). *dgcM* and *dgcQ* expression produced 2 proteins each, corresponding to their premature stop codon and adjacent methionine (Fig 6B), presumably translating the GGDEF domain in second reading frame (Table 1). *dgcM* expression also produced a third band at ∼15KDa, which does not match any reading frames. *dgcN* expression produced two bands at ∼17 KDa and ∼40 KDa. The 17 kDa size corresponds to the first N-terminal reading frame; however, the 40 KDa size corresponds with full-length DgcN. *dgcE* expression produced two bands at ∼45 KDa and 15 KDa. The 45 KDa corresponds to the N-terminal reading frame; however, the other band does not correspond with any potential *dgcE* reading frames. *E. coli* DgcE undergoes post-translational proteolysis, thus it’s possible that this processing is why the sizes we observed here did not match the predicted sizes (34, 49). *pdeR* expression also produced two bands; the first band at ∼18Kda corresponds to the first reading frame, however the second band at ∼22 KDa does not match any predicted reading frames.

### *dgcE* deletion reduces *S. flexneri* c-di-GMP levels

As expression of *S. flexneri dgcE, dgcQ* and *dgcN* pseudogenes in *E. coli* BL-21 significantly increased c-di-GMP levels, we hypothesized that degenerate DGCs also contribute to c-di-GMP production in *S. flexneri*. To test this hypothesis, we first deleted *dgcE* from the *S. flexneri* Δ4DGC background and generated a Δ5DGC strain (Δ4DGC+Δ*dgcE*). Then, we deleted *dgcQ* in the *S. flexneri* Δ5DGC background, to generate Δ6DGC (Δ4DGC+Δ*dgcE+*Δ*dgcQ*). We quantified the c-di-GMP levels of each of these strains using LC-MS/MS (Fig. 7). The *S. flexneri* Δ5DGC strain exhibited significantly reduced c-di-GMP levels compared to the WT *S. flexneri* strain; however, our Δ6DGC strain showed no significant difference compared to WT *S. flexneri* (Fig 7A). Notably, we still detected a c-di-GMP peak in the *S. flexneri* Δ6DGC strain, indicating that *dgcN* (or other DGCs) may also produce c-di-GMP (Fig. S4). We confirmed our findings using the riboswitch reporter and fluorescence microscopy. Measurements using the riboswitch reporter mirrored those taken by LC-MS/MS, and we observed significant reductions in c-di-GMP in both the Δ5DGC and Δ6DGC strains compared to the WT *S. flexneri* strain (Fig 7B).

**Figure 7.**
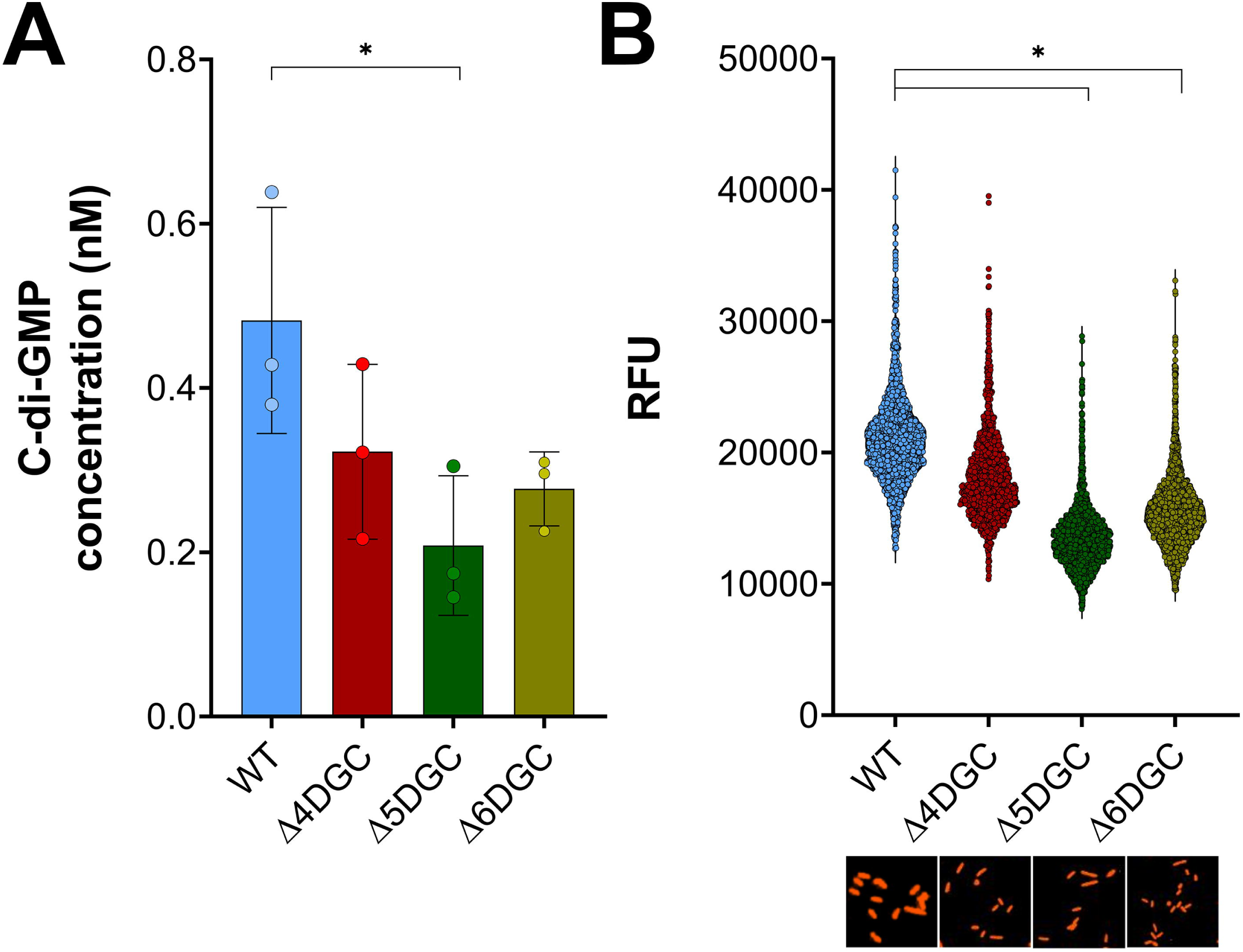
*S. flexneri* Δ5DGC and Δ6DGC strains have significantly reduced c-di-GMP levels. (A) *S. flexneri* Δ5DGC (Δ4DGC + Δ*dgcE*) and Δ6DGC (Δ4DGC + Δ*dgcE* + Δ*dgcQ*) c-di-GMP levels were compared to WT *S. flexneri* using LC-MS/MS. The *S. flexneri* Δ5DGC strain showed significantly lower c-di-GMP levels compared to the WT *S. flexneri* strain. In the *S. flexneri* Δ6DGC strain, c-di-GMP levels were reduced but not statistically significant than in the WT strain. (B) *S. flexneri* Δ5DGC and Δ6DGC strain c-di-GMP levels quantified using the c-di-GMP riboswitch reporter. C-di-GMP levels were significantly lower for the Δ5DGC or Δ6DGC strain in comparison to WT *S. flexneri*. Each symbol represents individual fluorescent cells for each strain. Images below the graph are individual fluorescent cells for each strain. Asterisks represent significant differences among strains as compared using one-way ANOVA with Sidak’s multiple comparisons posttest (*p<0.05*), error bars indicate standard deviation.

## Discussion

Because of the evolutionary relationship between commensal *E. coli* and pathogenic *S. flexneri*, we can use these two organisms as a model to examine how c-di-GMP signaling changes in the process of pathoadaptation (50). Only the four DGCs most conserved in other *Shigella* spp. (Fig. S3) have been retained with high fidelity in *S. flexneri* 2457T. One possible reason that these enzymes were conserved over others is signaling specificity, where specific DGCs alter c-di-GMP synthesis and associated phenotypes (31, 33, 34). Our data support this hypothesis, as there is a poor correlation between the expression of each of the four intact *S. flexneri* DGCs and the phenotypes we measured in this study. For example, *dgcF* expression produces less c-di-GMP than *dgcI* expression in *S. flexneri*, but it is the only DGC that significantly decreases acid sensitivity in AR1-inducing conditions. Our observation that *S. flexneri* GG→AA mutated *dgcI* regulated invasion and plaque phenotypes irrespective of catalytic activity is also suggestive that protein-protein interactions are enabling specific phenotypes (24, 51, 52). An alternative explanation regarding the discrepancy between measured c-di-GMP and various phenotypes resulting from DGC expression is that environmental regulation is contributing to differences we observe between c-di-GMP measurements and phenotypes, as environmental conditions in phenotypic assays do not precisely match those used for c-di-GMP measurements, and DGCs recognize environmental signals via their variable N-terminal domains that alter enzymatic activity at the C-terminal domain (33, 53).

We observed that c-di-GMP levels inside host cells were dynamic during infection (Fig. 3), and that *S. flexneri* DGCs produced c-di-GMP differentially in this environment (Fig. 4). *S. flexneri* virulence gene expression is dynamic during growth in an epithelial cell, and the timing correlates with the increase in c-di-GMP we observed (10, 51, 54–56). Expression of VCA0956 in *S. flexneri*, which raises c-di-GMP, significantly downregulates the expression of virulence genes, including those of the T3SS (20), and a similar phenomenon was also reported in *S. typhimurium* (24). Thus, it is possible that c-di-GMP is contributing to differential gene expression during growth in a host cell, but the signal transduction pathway linking c-di-GMP signaling and virulence gene regulation is still unknown.

Surprisingly, we found that deleting all four intact DGCs from the *S. flexneri* genome reduced c-di-GMP levels but did not eliminate c-di-GMP, and that expression of *S. flexneri* degenerate DGCs *dgcE*, *dgcQ*, and *dgcN* increased c-di-GMP (Fig 6,7A). The degenerate DGCs in *S. flexneri* are among those that synthesize c-di-GMP in other *Escherichia*/*Shigella* spp. (21, 57). Expression of these *S. flexneri* degenerate DGC genes results in multiple protein isoforms, some of which could potentially contain the GGDEF domain (Fig. 7B). *E. coli dgcE* is a large protein, encoding 10 transmembrane domains within the first 300 residues. It also encodes a predicted MASE1 domain and three PAS domains before a GGDEF and degenerate EAL domain. *E. coli* DgcE undergoes rapid proteolysis through an unknown protease, resulting in protein degradation. However, this pattern was not observed in the ΔMASE (N-terminal) DgcE, suggestive of proteolysis initiating from the N-terminal domain to the C-terminal domain (34, 49). DgcE initiates the expression of biofilm regulator CsgD and mediates curli and cellulose production in *E. coli*, facilitating exposure to host immune system (58–61). *S. flexneri dgcE* contains IS110 family transposase insertion, resulting in a premature stop codon and a frame shift; nevertheless, we found that deletion of *dgcE* from the Δ4DGC background significantly reduced c-di-GMP. *dgcE* has been annotated as degenerate in nine other pathogenic *E. coli* strains, including EHEC, EPEC, and EAEC isolates, suggesting a broader evolutionary adaptation associated with pathogenic host colonization (62).

In the case of *dgcQ* and *dgcN*, expression of these genes corresponded with two proteins each, one matching the size of the N-terminal reading frame and the other the full-length protein if it had no stop codon. It is unclear how *S. flexneri* is producing a full-length protein when both of these genes have nonsense mutations. One possibility is translational readthrough (63–67), or alternatively the bacteria could be inserting selenocysteine into these proteins in lieu of the stop codon. *dgcQ* mutations have been observed in at least six different strains of EAEC in the same specific pattern observed in *S. flexneri* 2457T, with an insertion of stop codon at 312 amino acid position and adjacent methionine (AUG) at 314 codon; this suggests that, like *dgcE*, the *dgcQ* adaptation is not unique to this specific *Shigella* strain (62).

*S. flexneri dgcM* expression in *E. coli* reduced c-di-GMP levels and was also expressed in two sizes, but these sizes corresponded to the N-terminal and C-terminal portions being separately translated. If one of these two proteins is the DgcM C-terminal portion, it is not clear how they are expressed without a ribosomal binding site. While *dgcM* does not contain a canonical Shine Dalgarno motif prior to the second start codon, it does contain an A-rich sequence upstream of the second reading frame (AAATAAAT), which can be sufficient to initiate translation (68). Alternatively, *dgcM* could be translating the second reading frame through translational coupling, where the ribosome translates an adjacent gene, modulated by the gene upstream to it (69, 70).

The mutations in *S. flexneri dgcQ*, *dgcE* and other degenerate DGCs exhibit a very similar pattern, suggesting that this type of mutation is a wider evolutionary mechanism to alter the activity of this enzyme. Deletion of dgcE and dgcQ from the *S. flexneri* Δ4DGC strain still did not eliminate c-di-GMP in the cell, indicating that other DGCs are still synthesizing the second messenger. It is possible that dgcN or dgcM are capable of c-di-GMP synthesis when expressed in *S. flexneri*, or the *pdeR* gene (containing both an EAL and GGDEF domain) has a premature nonsense mutation preceding a GGDEF domain, similar to *dgcE* and *dgcQ*.

## Experimental Procedures

### Strain construction and plasmids

*Shigella* strains used in this study were routinely cultured on tryptic soy broth agar (TSBA) with Congo red, and red colonies were selected to maintain the virulence plasmid (71). Antibiotic concentrations used were as follows until unless otherwise specified: 25 mg/ml ampicillin, 50 mg/ml chloramphenicol, and 20 mg/ml gentamicin, 50 mg/ml kanamycin, IPTG was added at 100μM. Other details of strains used are summarized in Table S1. *S. flexneri* Δ4DGC, Δ5DGC and Δ6DGC mutant strains were generated using homologous recombination (72). Briefly, single genes were deleted by insertion of the *cam* gene flanked by FRT sites, which were later removed by expressing flippase. DGC expression plasmids were constructed by restriction cloning, using the IPTG inducible pEVS143 (73) as the parental plasmid. The GG→AA mutants were generated by site-directed mutagenesis, using Phusion polymerase (NEB). All strains and plasmids were verified using sequencing. All primers used for homologous recombination and plasmid cloning are listed in Table S2. For infection assays, we used human Henle-407 cells (ATCC CCL-6, HeLa contamination line) as previously described (39).

### C-di-GMP quantification using LC-MS/MS method

Overnight cultures from single colonies were grown in LB media with appropriate antibiotics at 30°C with shaking. Next day, the overnight cultures were subcultured 1:100 in freshly prepared M63 media with Kan and IPTG. The optical density (OD_600_) of the subcultures grown at 37°C for ∼3 hours was recorded. From this point, the cultures were centrifuged at 4°C for 5 minutes. The supernatant was removed and the pellet was resuspended in 500 μl ice cold extraction buffer (40% Acetonitrile, 40% Methanol, and 0.1N formic acid). The resuspended pellet was incubated at −20°C for 30 minutes and then centrifuged for 20 minutes at 4°C. After centrifugation, the liquid was collected and stored at −80°C until analysis. Prior to analysis, samples were dried using a vacuum concentrator and then rehydrated in 100 μl water. Quantification of c-di-GMP was performed using a Quattro Premier XE mass spectrometer (Waters) coupled with an Acquity high performance liquid chromatography system (Waters) at the Western Michigan University Homer Stryker M.D. School of Medicine (WMeD), at mass 690.69→344.3. Using microscopy, we estimated the mean volume (∼2.36X10^15^) of *S. flexneri* cells which was used to determine the intracellular c-di-GMP. The intracellular c-di-GMP concentrations was determined by dividing the c-di-GMP extracted for each strain by the estimated mean volume. Total cell volume of extracted bacteria was calculated by multiplying the CFU/ml to the volume of one bacterium. Experiments were performed in three individual replicates. Graphs were plotted using Graphpad prism.

### Microscopy for c-di-GMP quantification using riboswitch reporter

*S. flexneri* strains containing DGC expression plasmids and the biosensor plasmid were grown in LB media with Kan and Amp at 30°C for ∼16 hours. Next day, the strains were subcultured in M63 media with Kan and Amp and were grown for ∼3.5 hours. After incubation, the cultures were diluted 1:1000 in M63 media with Kan, Amp and IPTG. Immediately, 100 μl of this suspension was transferred to a glass bottom 96 well plate, centrifuged for 10 minutes at 1000 x g, and cells were imaged using a BioTek Lionheart FX automated fluorescence microscope for 10 hours, with half hour interval. Samples were incubated at 37 °C during imaging. Time zero indicates the first microscopy read after adding IPTG and 10 minutes centrifugation. Our positive control, the VCA0956 expression strain, was used to optimize the Tetramethylrhodamine (TRITC) channel exposure. We used autofocus with a 25 μM Z stack projection and beacons were defined for each well. Fluorescence quantitation was performed using BioTek Gen5 software, using fluorescence autothreshold and size threshold between 2μM to 20 μM. An average cell count for each strain was ∼400 cells at time 0 and ∼4000 cells at the tenth hour for each strain. Experimental quantitation of fluorescence was performed three independent times.

### Host cell c-di-GMP kinetics read with microscopy

Henle-407 cells were split in a glass bottom 96 well plate 2 days before experiment to gain ∼80% confluency (1:3 dilution). One day before experiment, *S. flexneri* cultures were grown with Kan and Amp in LB media at 30°C for ∼16 hours. On the day of experiment, all the strains were sub cultured in M63 media with Kan, Amp and DOC (0.04% v/v) at 37°C for ∼4 hours. The OD_650_ was measured, and the cells were normalized to 2X10^9^ CFU/ml before infection. Infection was performed as previously described (39). Briefly, the normalized cells were used to infect the Henle-407 cells and centrifuged at 105 x g for 10 minutes. After centrifugation, the 96 well plate was incubated at 37°C with 5% CO_2_ for 30 minutes. After 30 minutes, the non-adherent population was removed by washing four times with PBS. After washing, 100 μl fresh tissue culture media (FluoroBrite DMEM; Thermo Fisher A1896701) with Kan, Amp, Gen, IPTG, 10mM Hepes and 1:1000 dilution Hoechst was added to each well and was immediately imaged for 10 hours with half an hour interval. Samples were incubated at 37 °C during imaging. Our positive control VCA0956 was used to setup the exposure settings for Hoechst (blue channel), the TRITC (red channel), and the phase contrast. Settings for analysis and calculations were same as detailed above.

### Acid resistance (AR1 and AR2)

Acid resistance was setup as previously described with modifications (41, 44). *S. flexneri* strains were grown overnight in LB media with Kan at 30°C for ∼16 hours. Next day, for AR1 system the strains were subcultured 1:100 in LB media with 50mM morpholineethanesulfonic acid (MES), Kan and IPTG at pH 5.5 overnight at 37°C with shaking. For AR2, strains were subcultured in LB media with 50mM MES, 0.4% glucose Kan and IPTG at pH 5.5 overnight at 37°C with no shaking. The following day, the OD_650_ was measured for all the strains (AR1 and AR2) and the bacterial cells were normalized to 5X10^8^ CFU/ml. The normalized cells were acid shocked in M9 media at pH 2.5 at 37°C (AR1) and M9 media at pH 2.5 with 50 mM glutamate at 37°C (AR2). As a control, each strain was also incubated in M9 media at pH 7. After one hour, cells were plated with serial dilution. Colony counts and the dilution factor was recorded to calculate the percent survival, determined by dividing the CFU/mL of acid shocked strains by the CFU/mL of the same strains incubated in pH 7 medium.

### Cell culture assays

The invasion assay was set up as previously described (39). Briefly, one day before experiment, Henle-407 were split in 6 well plates for ∼10% confluency. *S. flexneri* DGC strains were inoculated in LB media with Kan and incubated overnight at 30°C. Each strain was subcultured 1:100 in LB media with Kan, IPTG and 0.04% DOC for ∼4 hours. The OD_600_ was normalized to 2×10^9^ CFU/ml and 100 μl of suspension was added to individual wells to infect Henle cells. Plates were centrifuges for 10 minutes at 1000 rpm and then incubated for 30 minutes at 37°C with 5% CO_2_. Each well was washed four times with PBS and incubated again with Gentamycin for 45 minutes. After washing the cells twice with PBS, each well was stained with Giemsa stain for 5 minutes and washed with water to remove excess stain. Later, 300 cells were counted for positively invaded cells with 3 or more bacterial cells.

Plaque assay was setup using previously described protocols (39). Briefly, two days before experiment, Henle-407 cells were split in 6 wells plates for ∼80% confluency. Overnight grown strains were subcultured 1:100 in LB with Kan for ∼4 hours. The cells were normalized to 5X10^4^ CFU/ml and 100 μl was added to individual wells. Each 6 well plate was centrifuged and incubated for 45 minutes at 37°C with 5% CO_2_. Each well was washed four times with PBS and incubated with Kan, IPTG and 20% glucose for 48 hours. After incubation, wells were washed twice with PBS and stained with crystal violet for 2 minutes. All plates were later imaged and plaque size was measured using ImageJ software.

### Immunoblotting

Immunoblots were performed as previously described (20). Bacterial cultures were grown in LB broth at 37 °C. When the cells reached the mid-log phase of growth, the OD600 was read, and cells were collected by centrifugation and resuspended in SDS sample buffer (5% β-mercaptoethanol, 3% (wt/vol) SDS, 10% glycerol, 0.02% bromophenol blue, 63 mM Tris-Cl, pH 6.8) at a concentration of 2 x 10^9^ CFU / mL or 4 x 10^9^ CFU / mL (for DgcN and DgcE only). Samples were boiled for 10 min, and then electrophoresed in a 20% SDS-PAGE gel. After electrophoresis, proteins were transferred to a 0.45-μm-pore-size nitrocellulose membrane (Hybond-ECL; GE Healthcare) and incubated with a α-HIS antibody (Genscript A00186), followed by HRP-conjugated α-mouse antibody (Abclonal AS003). Signals were detected by developing the blot with a Pierce ECL detection kit (Thermo-Fisher Scientific).

### Data availability

We confirm that the data supporting the findings of this study are available within the article and its supplementary materials, and raw data are available from the corresponding author (BK) upon request.

## Acknowledgements

We would like to thank Dr. Chris Waters and Amber Bedore (Michigan State University) for providing us with the riboswitch reporter. We are also very thankful to Dr. Kevin Blair (WMU) and Paul Moorman (WMed) for their help with LC-MS/MS.

## Author contributions

R.O. contributed to the conceptualization, formal analysis, investigation, visualization, and writing the original draft. S.K. contributed to the investigation. P.J. contributed to methodology and resources. B.J.K. contributed to conceptualization, data curation, project administration, resources, supervision, and funding acquisition. All authors contributed to the review and editing of the final draft.

## Funding and additional information

This work is supported by startup funds and a grant from the Faculty Research and Creative Activities Award from Western Michigan University. Work in the lab was supported by the NIH R03 grant AI156432-01A1 (B.J.K.). The content is solely the responsibility of the authors and does not necessarily represent the official views of the National Institutes of Health.

## Abbreviations

c-di-GMP: cyclic-di-guanosine monophosphate
DGC: diguanylate cyclase
T3SS: type III secretion system
PDE: phosphodiesterase
IPTG: isopropyl-b-D-thiogalactopyranoside
RFU: relative fluorescence unit
AR: acid resistance
HPI: hours post infection
TSBA: tryptic soy broth agar
LB: Luria Broth

---

**Table S1.** Strains and plasmids used for this study.

| Strain | Parent Strain | Other information | Method for chromosomal mutation | Reference |
| --- | --- | --- | --- | --- |
| *Shigella flexneri* 2457T |  |  |  |  |
| Δ4DGC *S. flexneri* | *S. flexneri* 2457T | Deletion of *dgcC*, *dgcF*, *dgcl*, and *dgcP* | Homologous Recombination | |
| Δ5DGC *S. flexneri* | Δ4DGC *S. flexneri* | Deletion of *dgcC*, *dgcF*, *dgcl*, *dgcP*, and *dgcE* | Homologous Recombination | |
| Δ6DGC *S. flexneri* | Δ5DGC *S. flexneri* | Deletion of *dgcC*, *dgcF*, *dgcl*, *dgcP*, *dgcE*, and *dgcQ* | Homologous Recombination | |
| *E. coli* BL21 |  |  |  |  |
| **Plasmids** |  |  |  |  |
| Plasmid | Parent Plasmid | Other information | Antibiotic marker plasmid | Reference |
| *dgcC* expression | pEVS143 | pTac promotor (IPTG inducible) + Sf *dgcC* | Kanamycin |  |
| *dgcF* expression | pEVS143 | pTac promotor (IPTG inducible) + Sf *dgcF* | Kanamycin |  |
| *dgcl* expression | pEVS143 | pTac promotor (IPTG inducible) + Sf *dgcl* | Kanamycin |  |
| *dgcP* expression | pEVS143 | pTac promotor (IPTG inducible) + Sf *dgcP* | Kanamycin |  |
| GG→AA *dgcC* | pEVS143 | pTac promotor (IPTG inducible) + Sf *dgcC* GG→AA | Kanamycin |  |
| GG→AA *dgcF* | pEVS143 | pTac promotor (IPTG inducible) + Sf *dgcF*→GGAA | Kanamycin |  |
| GG→AA *dgcl* | pEVS143 | pTac promotor (IPTG inducible) + Sf *dgcl* GG→AA | Kanamycin |  |
| GG→AA *dgcP* | pEVS143 | pTac promotor (IPTG inducible) + Sf *dgcP* GG→AA | Kanamycin |  |
| Riboswitch Reporter | RSF1010 |  | Ampicillin | (38) |
| *dgcM* expression | pET 28a(+) | T7 Promotor (IPTG inducible) | Kanamycin |  |
| *dgcN* expression | pET 28a(+) | T7 Promotor (IPTG inducible) | Kanamycin |  |
| *dgcE* expression | pET 28a(+) | T7 Promotor (IPTG inducible) | Kanamycin |  |
| *dgcQ* expression | pET 28a(+) | T7 Promotor (IPTG inducible) | Kanamycin |  |
| *dgcO* expression | pET 28a(+) | T7 Promotor (IPTG inducible) | Kanamycin |  |
| *dgcJ* expression | pET 28a(+) | T7 Promotor (IPTG inducible) | Kanamycin |  |
| *pdeR* expression | pET 28a(+) | T7 Promotor (IPTG inducible) | Kanamycin |  |

**Table S2.**
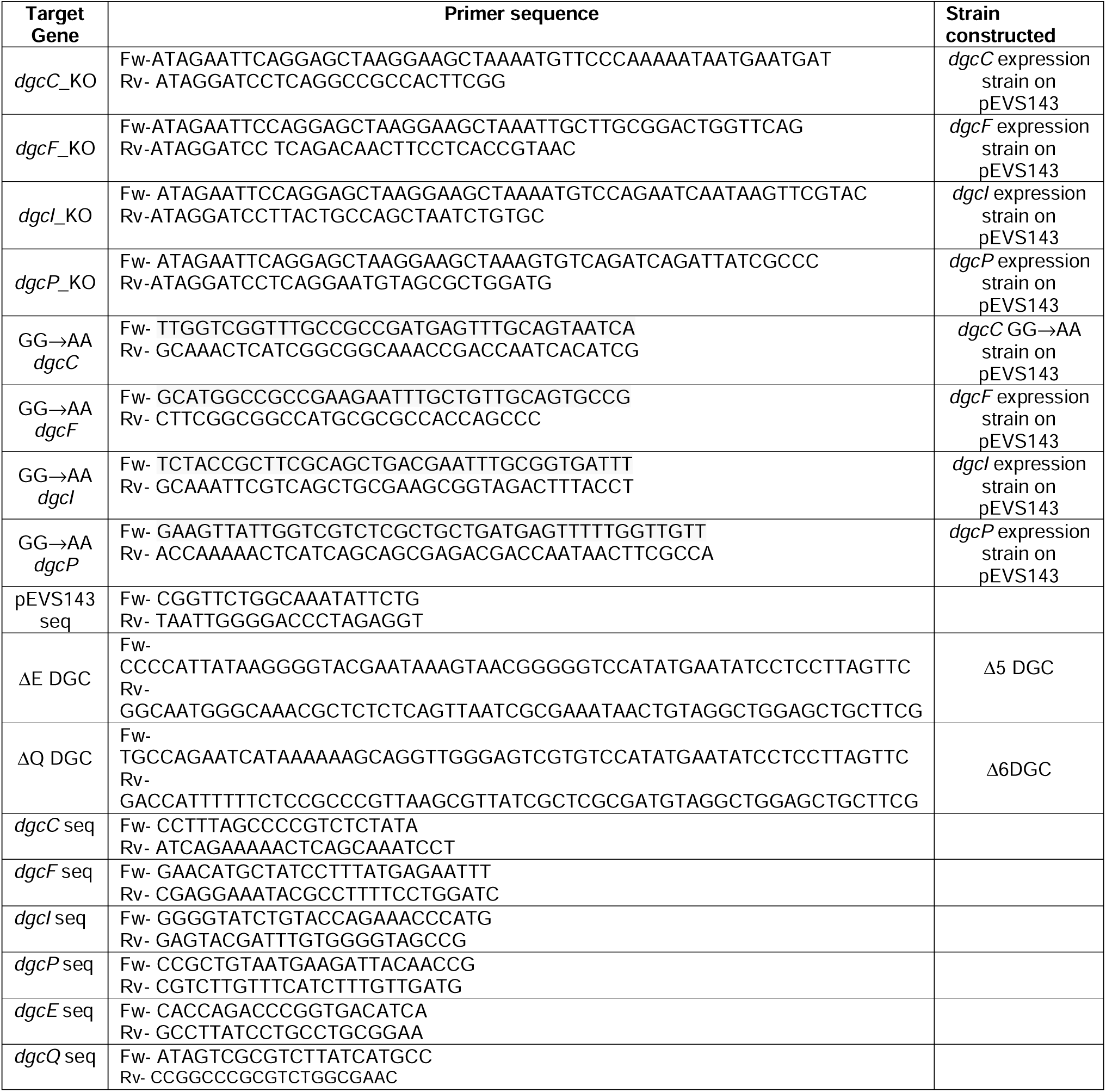
Primers used in this study.

**Supplementary Figure 1**

*S. flexneri* DGC-expressing strains showed no growth defects. Individual *S. flexneri* DGCs and the GG→AA mutant DGCs expressed on a Δ4DGC background with kanamycin and IPTG grew comparably to the Δ4DGC strain. Optical density (OD_600_) was measured every hour. Symbols represent the mean of three technical replicates for each strain and error bars represents standard deviation. There were no significant differences in growth between strains as analyzed by two-way ANOVA with Sidak’s multiple comparisons posttest.

**Supplementary Figure 2**

C-di-GMP-dependent fluorescence of the *S. flexneri* Δ4DGC strain expressing (A) *dgcC*, (B) *dgcF*, (C) *dgcI*, or (D) *dgcP*, alongside their respective GG→AA mutant alleles was quantified using a platereader. We found that the fluorescence, as quantified by platereader, followed the same patterns as fluorescence quantified by microscopy. Symbols indicate the mean and error bars the standard deviation of three independent experiments.

**Supplementary Figure 3.**

Four intact DGCs found in *S. flexneri* are highly conserved DGCs. We compiled a database of *Shigella* protein sequences derived from NCBI (TaxID:620), filtering for complete and annotated genomes, resulting in protein sequences from 219 strains. We then queried this database (BLAST) using each DGC protein sequence from *E. coli* K12 substr. MG1655 (Accession: NC_000913). We found that the 4 intact *S. flexneri* 2457T DGCs were the most conserved in other *Shigella* spp, while degenerate DGCs were present in a lower percentage of *Shigella* spp. DGCs are arranged based on the percent conservation, starting from the highly conserved DGC to the lowest conserved (left to right). Here, intact DGCs represent *S. flexneri* 2457T DGCs that are annotated as functional. Degenerate DGCs represent genes that are annotated as non-functional based on the mutations.

**Supplementary Figure 4.**

LC-MS/MS spectra showing c-di-GMP peaks for *S. flexneri* WT, Δ4DGC, Δ5DGC and Δ6DGC. The c-di-GMP peak (highlighted in gray) was highest for WT *S. flexneri*. Spectra were arranged using Adobe Photoshop; image transformations included only perspective transformation and contrast adjustment were applied identically to each spectrum panel.

## References

1. Kotloff KL, Winickoff JP, Ivanoff B, Clemens JD, Swerdlow DL, Sansonetti PJ, Adak GK, Levine MM. 1999. Global burden of Shigella infections: implications for vaccine development and implementation of control strategies. Bull World Health Organ 77:651–666.

2. Bennish ML, Wojtyniak BJ. 1991. Mortality Due to Shigellosis: Community and Hospital Data. Clin Infect Dis 13:S245–251.

3. Mantis N, Prévost MC, Sansonetti P. 1996. Analysis of epithelial cell stress response during infection by Shigella flexneri. Infect Immun 64:2474–2482.

4. Maurelli AT, Blackmon B, Curtiss R. 1984. Temperature-dependent expression of virulence genes in Shigella species. Infect Immun 43:195–201.

5. CDC. 2013. Antibiotic Resistance Threats in the United States. U.S. Department of Health and Human Services, Atlanta, GA.

6. CDC. 2023. Increase in Extensively Drug-Resistant Shigellosis in the United States. Health Alert Network. CDCHAN-00486.

7. Jennison AV, Verma NK. 2004. *Shigella flexneri* infection: pathogenesis and vaccine development. FEMS Microbiol Rev 28:43–58.

8. Dorman CJ. 2009. The Virulence Plasmids of Shigella flexneri, p. 151–170. *In* Schwartz, E (ed.), Microbial Megaplasmids. Springer Berlin Heidelberg, Berlin, Heidelberg.

9. Gall TL, Mavris M, Martino MC, Bernardini ML, Denamur E, Parsot C. 2005. Analysis of virulence plasmid gene expression defines three classes of effectors in the type III secretion system of *Shigella flexneri*. Microbiology 151:951–962.

10. Blocker A, Gounon P, Larquet E, Niebuhr K, Cabiaux V, Parsot C, Sansonetti P. 1999. The tripartite type III secreton of *Shigella flexneri* inserts IpaB and IpaC into host membranes. 3. J Cell Biol 147:683–693.

11. Suzuki T, Sasakawa C. 2001. Molecular basis of the intracellular spreading of *Shigella*. 10. Infect Immun 69:5959–66.

12. Zychlinsky A, Prevost MC, Sansonetti PJ. 1992. *Shigella-flexneri* induces apoptosis in infected macrophages. 6382. Nature 358:167–169.

13. Goldberg MB, Theriot JA. 1995. *Shigella flexneri* surface protein IcsA is sufficient to direct actin-based motility. Proc Natl Acad Sci U S A 92:6572–6576.

14. Chiang I-L, Wang Y, Fujii S, Muegge BD, Lu Q, Tarr PI, Stappenbeck TS. 2021. Biofilm Formation and Virulence of *Shigella flexneri* Are Modulated by pH of Gastrointestinal Tract. Infect Immun 89:e00387–21.

15. Cheng F, Wang J, Peng J, Yang J, Fu H, Zhang X, Xue Y, Li W, Chu Y, Jin Q. 2007. Gene expression profiling of the pH response in *Shigella flexneri* 2a. FEMS Microbiol Lett 270:12–20.

16. Faherty CS, Redman JC, Rasko DA, Barry EM, Nataro JP. 2012. Shigella flexneri effectors OspE1 and OspE2 mediate induced adherence to the colonic epithelium following bile salts exposure: OspE1 and OspE2 enhance adherence to epithelial cells. Mol Microbiol 85:107–121.

17. Pope LM, Reed KE, Payne SM. 1995. Increased protein secretion and adherence to HeLa cells by *Shigella* spp. following growth in the presence of bile salts. 9. Infect Immun 63:3642–8.

18. Nickerson KP, Chanin RB, Sistrunk JR, Rasko DA, Fink PJ, Barry EM, Nataro JP, Faherty CS. 2017. Analysis of Shigella flexneri Resistance, Biofilm Formation, and Transcriptional Profile in Response to Bile Salts. Infect Immun 85:e01067–16, e01067-16.

19. Koestler BJ, Fisher CR, Payne SM. 2018. Formate Promotes *Shigella* Intercellular Spread and Virulence Gene Expression. mBio 9:e01777–18, /mbio/9/5/mBio.01777-18.

20. Ojha R, Dittmar AA, Severin GB, Koestler BJ. 2021. *Shigella flexneri* Diguanylate Cyclases Regulate Virulence. J Bacteriol 203:e0024221.

21. Sen T, Verma NK. 2022. YfiB: An Outer Membrane Protein Involved in the Virulence of *Shigella flexneri*. Microorganisms 10:653.

22. Conner JG, Zamorano-Sánchez D, Park JH, Sondermann H, Yildiz FH. 2017. The ins and outs of cyclic di-GMP signaling in *Vibrio cholerae*. Curr Opin Microbiol 36:20–29.

23. D’Argenio DA. 2004. Cyclic di-GMP as a bacterial second messenger. Microbiology 150:2497–2502.

24. Ahmad I, Lamprokostopoulou A, Le Guyon S, Streck E, Barthel M, Peters V, Hardt WD, Romling U. 2011. Complex c-di-GMP signaling networks mediate the transition between biofilm formation and virulence properties in *Salmonella enterica* serovar Typhimurium. Int J Med Microbiol 301:84–85.

25. Beyhan S, Tischler AD, Camilli A, Yildiz FH. 2006. Transcriptome and phenotypic responses of *Vibrio cholerae* to increased cyclic di-GMP level. J Bacteriol 188:3600–3613.

26. Tischler AD, Camilli A. 2004. Cyclic diguanylate (c-di-GMP) regulates *Vibrio cholerae* biofilm formation: c-di-GMP regulates *V. cholerae* biofilm. Mol Microbiol 53:857–869.

27. Lamprokostopoulou A, Monteiro C, Rhen M, RÃ¶mling U. 2010. Cyclic di-GMP signalling controls virulence properties of *Salmonella enterica* serovar Typhimurium at the mucosal lining. Environ Microbiol 12:40–53.

28. Solano C, Garcia B, Latasa C, Toledo-Arana A, Zorraquino V, Valle J, Casals J, Pedroso E, Lasa I. 2009. Genetic reductionist approach for dissecting individual roles of GGDEF proteins within the c-di-GMP signaling network in *Salmonella*. 19. Proc Natl Acad Sci U S A 106:7997–8002.

29. Hunter JL, Severin GB, Koestler BJ, Waters CM. 2014. The *Vibrio cholerae* diguanylate cyclase VCA0965 has an AGDEF active site and synthesizes cyclic di-GMP. BMC Microbiol 14:22.

30. Koestler BJ, Waters CM. 2014. Bile Acids and Bicarbonate Inversely Regulate Intracellular Cyclic di-GMP in *Vibrio cholerae*. Infect Immun 82:3002–3014.

31. Kulasakara H, Lee V, Brencic A, Liberati N, Urbach J, Miyata S, Lee DG, Neely AN, Hyodo M, Hayakawa Y, Ausubel FM, Lory S. 2006. Analysis of *Pseudomonas aeruginosa* diguanylate cyclases and phosphodiesterases reveals a role for bis-(3 ’-5 ’)-cyclic-GMP in virulence. 8. Proc Natl Acad Sci U S A 103:2839–44.

32. Massie JP, Reynolds EL, Koestler BJ, Cong J-P, Agostoni M, Waters CM. 2012. Quantification of high-specificity cyclic diguanylate signaling. Proc Natl Acad Sci 109:12746–12751.

33. Waters CM. 2012. The Meteoric Rise of the Signaling Molecule Cyclic di-GMP: During the past decade, research on c-di-GMP expanded greatly, uncovering several roles it plays among bacteria. Microbe Mag 7:353–359.

34. Sarenko O, Klauck G, Wilke FM, Pfiffer V, Richter AM, Herbst S, Kaever V, Hengge R. 2017. More than Enzymes That Make or Break Cyclic Di-GMP—Local Signaling in the Interactome of GGDEF/EAL Domain Proteins of *Escherichia coli*. mBio 8:e01639–17, /mbio/8/5/e01639-17.

35. Hengge R, Galperin MY, Ghigo J-M, Gomelsky M, Green J, Hughes KT, Jenal U, Landini P. 2016. Systematic Nomenclature for GGDEF and EAL Domain-Containing Cyclic Di-GMP Turnover Proteins of Escherichia coli: TABLE 1. J Bacteriol 198:7–11.

36. Wei J, Goldberg MB, Burland V, Venkatesan MM, Deng W, Fournier G, Mayhew GF, Plunkett G, Rose DJ, Darling A, Mau B, Perna NT, Payne SM, Runyen-Janecky LJ, Zhou S, Schwartz DC, Blattner FR. 2003. Complete genome sequence and comparative genomics of *Shigella flexneri* serotype 2a strain 2457T. Infect Immun 71:2775–2786.

37. Da Re S, Ghigo J-M. 2006. A CsgD-Independent Pathway for Cellulose Production and Biofilm Formation in Escherichia coli. J Bacteriol 188:3073–3087.

38. Zhou H, Zheng C, Su J, Chen B, Fu Y, Xie Y, Tang Q, Chou S-H, He J. 2016. Characterization of a natural triple-tandem c-di-GMP riboswitch and application of the riboswitch-based dual-fluorescence reporter. Sci Rep 6:20871.

39. Koestler BJ, Ward CM, Payne SM. 2018. *Shigella* Pathogenesis Modeling with Tissue Culture Assays. Curr Protoc Microbiol 50:e57.

40. Lin J, Lee IS, Frey J, Slonczewski JL, Foster JW. 1995. Comparative analysis of extreme acid survival in Salmonella typhimurium, *Shigella flexneri*, and *Escherichia coli*. J Bacteriol 177:4097–4104.

41. Jennison AV, Verma NK. 2007. The acid-resistance pathways of *Shigella flexneri* 2457T. Microbiology 153:2593–2602.

42. Waterman SR, Small PLC. 2003. The glutamate-dependent acid resistance system of *Escherichia coli* and *Shigella flexneri* is inhibited in vitro by l-*trans* - pyrrolidine-2,4-dicarboxylic acid. FEMS Microbiol Lett 224:119–125.

43. Casalino M, Prosseda G, Barbagallo M, Iacobino A, Ceccarini P, Carmela Latella M, Nicoletti M, Colonna B. 2010. Interference of the CadC regulator in the arginine-dependent acid resistance system of Shigella and enteroinvasive *E. coli*. Int J Med Microbiol 300:289–295.

44. Weatherspoon-Griffin N, Wing HJ. 2016. Characterization of SlyA in *Shigella flexneri* Identifies a Novel Role in Virulence. Infect Immun 84:1073–1082.

45. Waterman SR, Small PLC. 1996. Identification of σ ^s^ -dependent genes associated with the stationary-phase acid-resistance phenotype of *Shigella flexneri*. Mol Microbiol 21:925–940.

46. Chanin RB, Nickerson KP, Llanos-Chea A, Sistrunk JR, Rasko DA, Kumar DKV, de la Parra J, Auclair JR, Ding J, Li K, Dogiparthi SK, Kusber BJD, Faherty CS. 2019. *Shigella flexneri* Adherence Factor Expression in *In Vivo* -Like Conditions. mSphere 4:e00751–19, /msphere/4/6/mSphere751-19.

47. Jenal U, Malone J. 2006. Mechanisms of cyclic-di-GMP signaling in bacteria. Annu Rev Genet 40:385–407.

48. Valentini M, Filloux A. 2016. Biofilms and Cyclic di-GMP (c-di-GMP) Signaling: Lessons from *Pseudomonas aeruginosa* and Other Bacteria. J Biol Chem 291:12547–12555.

49. Pfiffer V, Sarenko O, Possling A, Hengge R. 2019. Genetic dissection of Escherichia coli’s master diguanylate cyclase DgcE: Role of the N-terminal MASE1 domain and direct signal input from a GTPase partner system. PLOS Genet 15:e1008059.

50. Prosseda G, Di Martino ML, Campilongo R, Fioravanti R, Micheli G, Casalino M, Colonna B. 2012. Shedding of genes that interfere with the pathogenic lifestyle: the *Shigella* model. Res Microbiol 163:399–406.

51. Pieper R, Fisher CR, Suh M-J, Huang S-T, Parmar P, Payne SM. 2013. Analysis of the Proteome of Intracellular *Shigella flexneri* Reveals Pathways Important for Intracellular Growth. 12. Infect Immun 81:4635–4648.

52. Ahmad I, Cimdins A, Beske T, Römling U. 2017. Detailed analysis of c-di-GMP mediated regulation of csgD expression in Salmonella typhimurium. BMC Microbiol 17:27.

53. Dayton H, Smiley MK, Forouhar F, Harrison JJ, Price-Whelan A, Dietrich LEP. 2020. Sensory Domains That Control Cyclic di-GMP-Modulating Proteins: A Critical Frontier in Bacterial Signal Transduction, p. 137–158. In Chou, S-H, Guiliani, N, Lee, VT, Römling, U (eds.), Microbial Cyclic Di-Nucleotide Signaling. Springer International Publishing, Cham.

54. Ménard R, Sansonetti PJ, Parsot C. 1993. Nonpolar mutagenesis of the ipa genes defines IpaB, IpaC, and IpaD as effectors of Shigella flexneri entry into epithelial cells. J Bacteriol 175:5899–5906.

55. Bourdet-Sicard R, Rudiger M, Jockusch BM, Gounon P, Sansonetti PJ, Nhieu GT. 1999. Binding of the Shigella protein IpaA to vinculin induces F-actin depolymerization. 21. EMBO J 18:5853–62.

56. Headley VL, Payne SM. 1990. Differential protein expression by *Shigella flexneri* in intracellular and extracellular environments. 11. Proc Natl Acad Sci U S A 87:4179–83.

57. Serra DO, Hengge R. 2019. A c-di-GMP-Based Switch Controls Local Heterogeneity of Extracellular Matrix Synthesis which Is Crucial for Integrity and Morphogenesis of *Escherichia coli* Macrocolony Biofilms. J Mol Biol 431:4775– 4793.

58. Lindenberg S, Klauck G, Pesavento C, Klauck E, Hengge R. 2013. The EAL domain protein YciR acts as a trigger enzyme in a c-di-GMP signalling cascade in E. coli biofilm control. EMBO J 32:2001–2014.

59. Pesavento C, Becker G, Sommerfeldt N, Possling A, Tschowri N, Mehlis A, Hengge R. 2008. Inverse regulatory coordination of motility and curli-mediated adhesion in *Escherichia coli*. Genes Dev 22:2434–2446.

60. Sukupolvi S, Edelstein A, Rhen M, Normark SJ, Pfeifer JD. 1997. Development of a murine model of chronic *Salmonella* infection. Infect Immun 65:838–842.

61. Sakellaris H, Hannink NK, Rajakumar K, Bulach D, Hunt M, Sasakawa C, Adler B. 2000. Curli Loci of Shigella spp. Infect Immun 68:3780–3783.

62. Povolotsky TL, Hengge R. 2016. Genome-Based Comparison of Cyclic Di-GMP Signaling in Pathogenic and Commensal *Escherichia coli* Strains. J Bacteriol 198:111–126.

63. Dabrowski M, Bukowy-Bieryllo Z, Zietkiewicz E. 2015. Translational readthrough potential of natural termination codons in eucaryotes--The impact of RNA sequence. RNA Biol 12:950–958.

64. Sharma J, Du M, Wong E, Mutyam V, Li Y, Chen J, Wangen J, Thrasher K, Fu L, Peng N, Tang L, Liu K, Mathew B, Bostwick RJ, Augelli-Szafran CE, Bihler H, Liang F, Mahiou J, Saltz J, Rab A, Hong J, Sorscher EJ, Mendenhall EM, Coppola CJ, Keeling KM, Green R, Mense M, Suto MJ, Rowe SM, Bedwell DM. 2021. A small molecule that induces translational readthrough of CFTR nonsense mutations by eRF1 depletion. Nat Commun 12:4358.

65. Karki P, Carney TD, Maracci C, Yatsenko AS, Shcherbata HR, Rodnina MV. 2022. Tissue-specific regulation of translational readthrough tunes functions of the traffic jam transcription factor. Nucleic Acids Res 50:6001–6019.

66. Fan Y, Evans CR, Barber KW, Banerjee K, Weiss KJ, Margolin W, Igoshin OA, Rinehart J, Ling J. 2017. Heterogeneity of Stop Codon Readthrough in Single Bacterial Cells and Implications for Population Fitness. Mol Cell 67:826–836.e5.

67. Zhang H, Lyu Z, Fan Y, Evans CR, Barber KW, Banerjee K, Igoshin OA, Rinehart J, Ling J. 2020. Metabolic stress promotes stop-codon readthrough and phenotypic heterogeneity. Proc Natl Acad Sci 117:22167–22172.

68. Saito K, Green R, Buskirk AR. 2020. Translational initiation in E. coli occurs at the correct sites genome-wide in the absence of mRNA-rRNA base-pairing. eLife 9:e55002.

69. Levin-Karp A, Barenholz U, Bareia T, Dayagi M, Zelcbuch L, Antonovsky N, Noor E, Milo R. 2013. Quantifying Translational Coupling in *E. coli* Synthetic Operons Using RBS Modulation and Fluorescent Reporters. ACS Synth Biol 2:327–336.

70. Tian T, Salis HM. 2015. A predictive biophysical model of translational coupling to coordinate and control protein expression in bacterial operons. Nucleic Acids Res 43:7137–7151.

71. Payne SM. 2019. Laboratory Cultivation and Storage of *Shigella*. Curr Protoc Microbiol 55.

72. Datsenko KA, Wanner BL. 2000. One-step inactivation of chromosomal genes in Escherichia coli K-12 using PCR products. Proc Natl Acad Sci 97:6640–6645.

73. Dunn AK, Millikan DS, Adin DM, Bose JL, Stabb EV. 2006. New *rfp*- and pES213-derived tools for analyzing symbiotic *Vibrio fischeri* reveal patterns of infection and *lux* expression *in situ*. 1. Appl Environ Microbiol 72:802–810.

