## Supplementary figures and images for "Intact and Degenerate Diguanylate Cyclases regulate *Shigella* Cyclic di-GMP"

### Figure S1.tif

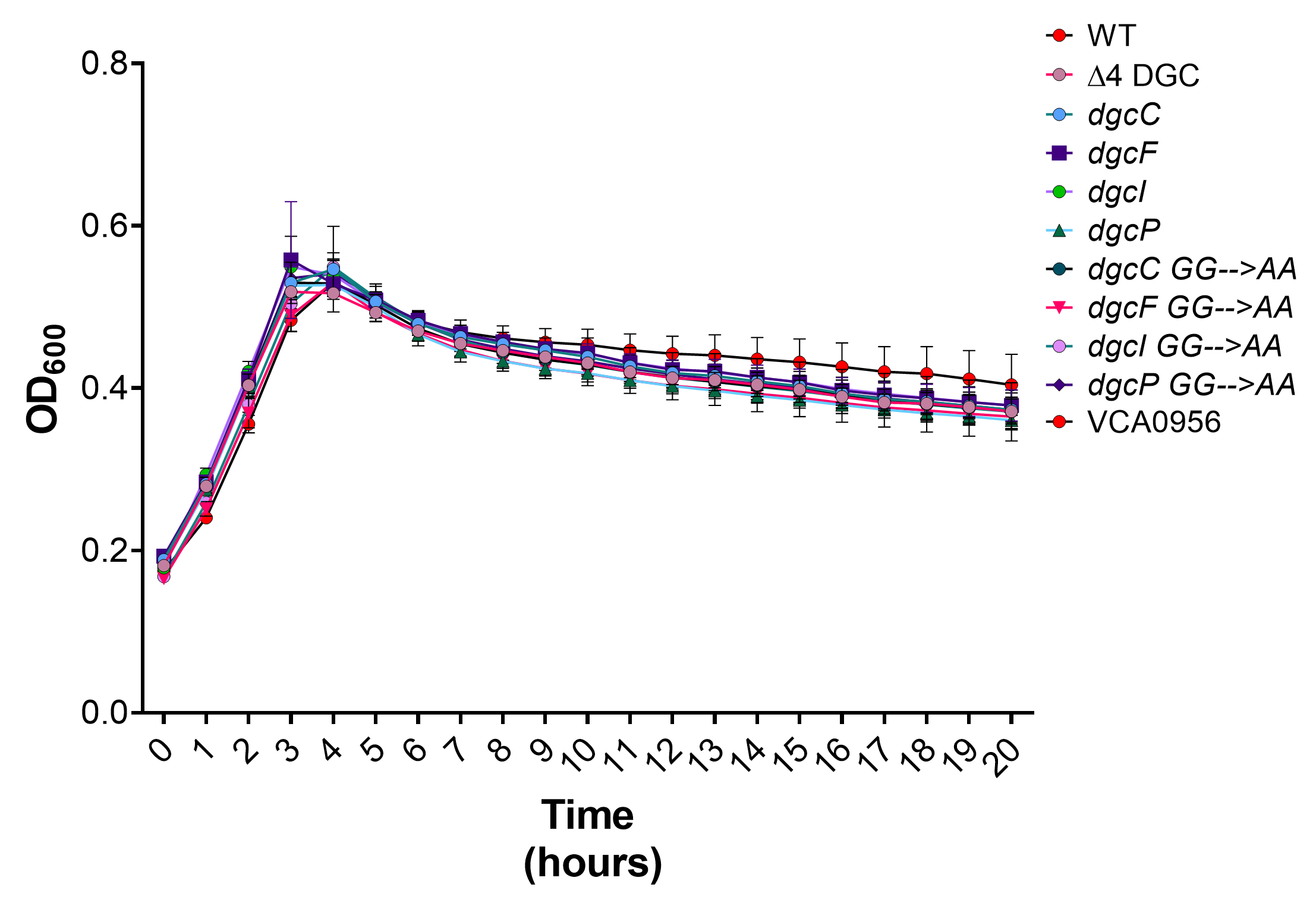

### Figure S2.tif

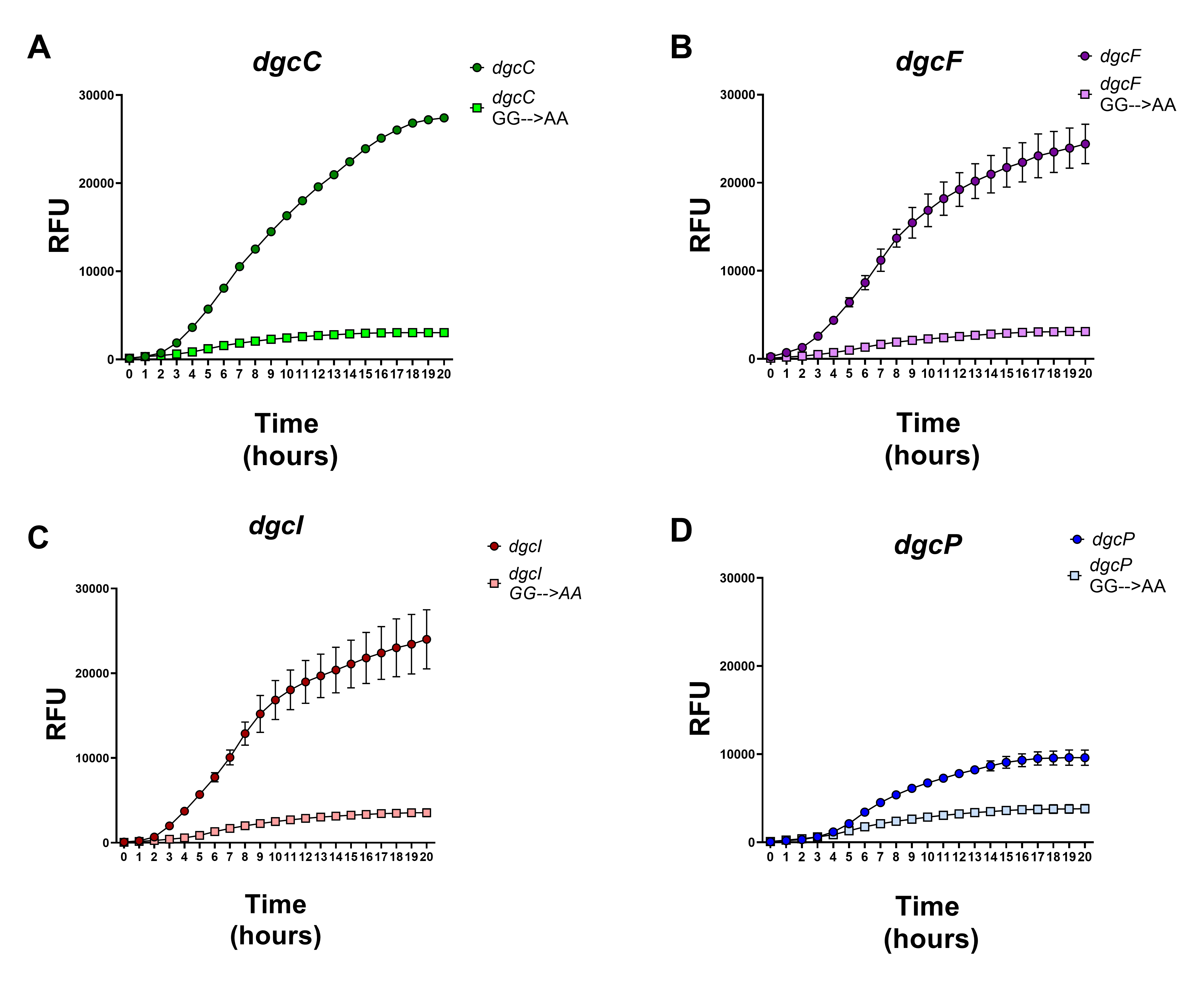

### Figure S3.tif

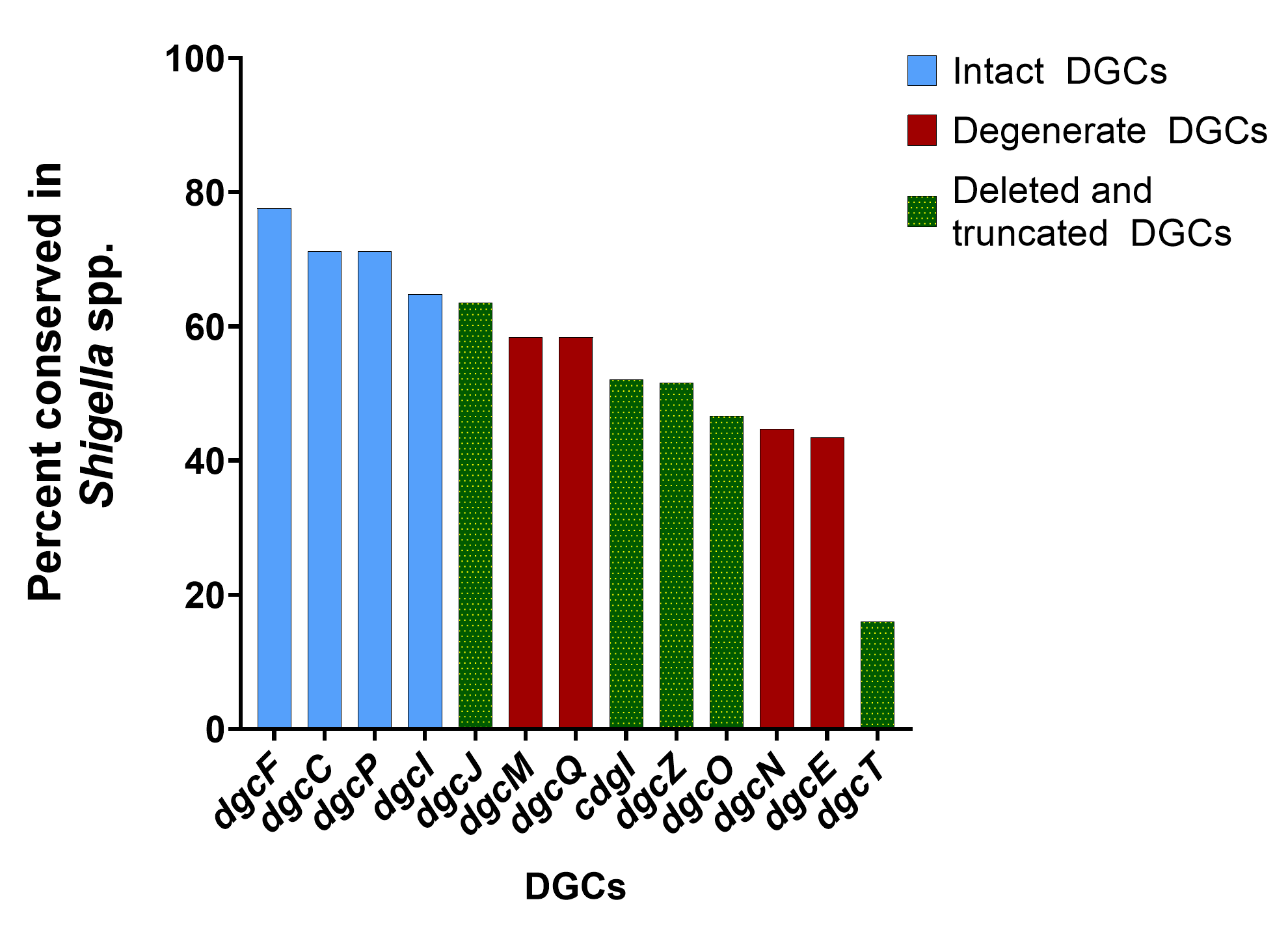

### Figure S4.tif

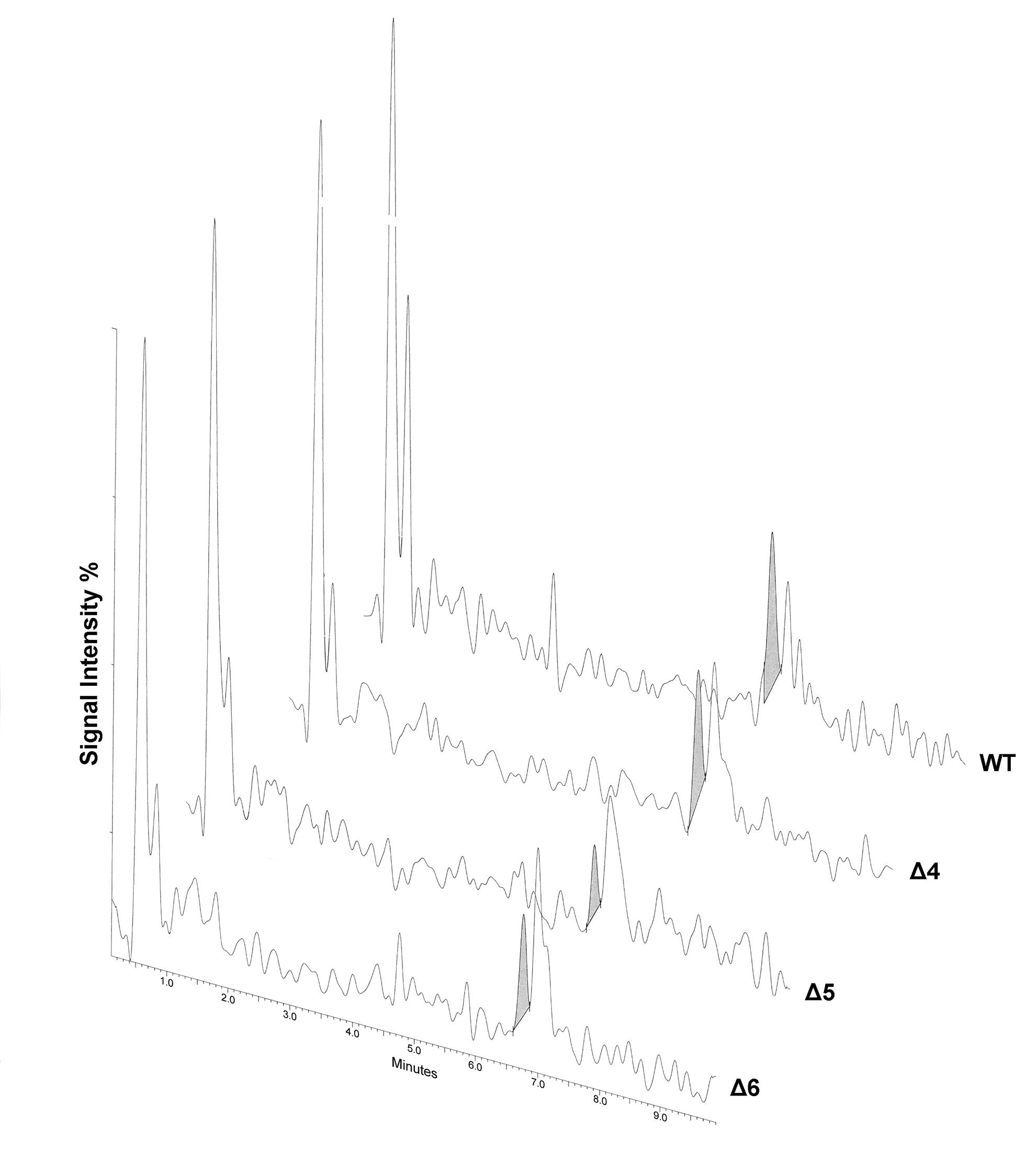
